# Quantitative profiling of intrinsic dCas9-DNA recognition reveals key determinants of guide RNA performance

**DOI:** 10.64898/2026.08.10.743836

**Authors:** Wei Zhu, Michael Tian, Yuncheng Duan, Samuel J. Reisman, Samantha E. Miller, Evan Corden, Maria ter Weele, Lingyun Song, Jameson Blount, Alexias Safi, Jacob Schreiber, Charles A. Gersbach, Gregory E. Crawford, Raluca Gordân

## Abstract

CRISPR technologies based on nuclease-deactivated Cas9 (dCas9) rely on programmable DNA binding rather than DNA cleavage, yet the intrinsic DNA-recognition properties that govern optimal guide RNA (gRNA) performance remain poorly understood. Existing approaches either measure genomic occupancy in cells or infer dCas9 behavior from cleavage-based Cas9 datasets, despite DNA binding being substantially more permissive than DNA cleavage. Here we introduce TANGO (Targeted Array-based Nucleic acid-Guided Occupancy), a high-density DNA-array platform that quantitatively profiles intrinsic dCas9:gRNA binding across tens of thousands of DNA targets in a cell-free system. TANGO captures established features of dCas9 target recognition, while providing substantially greater sensitivity than prior assays. Comparison with ChIP-seq data demonstrates that intrinsic DNA-binding specificity is a major driver of genomic occupancy and reveals that chromatin accessibility modulates the intrinsic binding affinity required for dCas9 recruitment. Across CRISPRi/a guides, TANGO identifies multiple independent biochemical determinants of guide performance—including on-target affinity, mismatch tolerance, and ribonucleoprotein assembly—and flags problematic and highly promiscuous guides overlooked by current specificity metrics. Unexpectedly, some guides retain substantial guide-directed DNA binding even in the absence of a protospacer-adjacent motif (PAM), revealing an additional dimension of dCas9 specificity. Together, these results establish intrinsic DNA recognition as a quantitative and experimentally accessible determinant of dCas9 function, providing a framework for improving guide selection and enhancing the precision of CRISPR technologies.

## INTRODUCTION

The CRISPR-Cas9 system has transformed genome engineering by enabling programmable recognition of virtually any DNA sequence through a guide RNA (gRNA). While nuclease-active Cas9 is widely used for genome editing, many of the fastest-growing CRISPR technologies instead employ catalytically deactivated Cas9 (dCas9), which retains sequence-specific DNA binding but lacks nuclease activity. By fusing dCas9 to transcriptional regulators, chromatin modifiers, base-editing enzymes, fluorescent proteins, or other functional domains, programmable DNA binding can be harnessed to modulate gene expression^1–9^, edit epigenetic states^10–13^, image genomic loci^14^, and perform numerous additional functions without introducing double-strand DNA breaks.

For these dCas9-based technologies, successful targeting depends fundamentally on DNA binding rather than DNA cleavage. Strong occupancy of the intended genomic locus is required for the attached effector to perform its function, while binding to unintended sites can reduce on-target activity or produce undesirable off-target effects. Despite the central importance of DNA binding, guide selection for CRISPR interference (CRISPRi), CRISPR activation (CRISPRa), and related technologies remains largely guided by computational models^15–18^ developed from cleavage-based datasets generated with nuclease-active Cas9^15,16,19^. However, DNA binding is substantially more permissive than DNA cleavage^20–22^, and many genomic sites that bind dCas9 are not detectably cleaved by Cas9^21–23^. Consequently, cleavage-based specificity measurements provide only an incomplete view of the DNA-recognition properties that ultimately determine dCas9 performance.

A major obstacle to understanding dCas9 specificity is that genomic occupancy measured in cells reflects both intrinsic DNA recognition by the dCas9:gRNA ribonucleoprotein (RNP) complex (i.e. recognition driven by the molecular properties of the RNP and the DNA sequence), and numerous extrinsic cellular factors including chromatin accessibility, DNA methylation, transcription factor occupancy, and local genome architecture. Genome-wide methods, such as ChIP-seq^23,24^ and CUT&Run/CUT&Tag^25–27^, therefore measure the combined outcome of intrinsic binding affinity and extrinsic cellular context, making it difficult to determine the exact causes of why some guides bind more efficiently than others. Conversely, existing high-throughput *in vitro* assays capable of measuring intrinsic dCas9 binding^28–31^ have remained limited in either sensitivity or in scale, restricting systematic characterization across diverse gRNAs and target sequences. As a result, the intrinsic biochemical properties governing dCas9-DNA recognition remain incompletely understood.

Here we introduce TANGO (Targeted Array-based Nucleic acid-Guided Occupancy), a high-density DNA-array platform that quantitatively measures intrinsic RNP binding across tens of thousands of designed DNA sequences in a cell-free system. By directly separating sequence-intrinsic DNA recognition from extrinsic chromatin-dependent effects, TANGO enables comprehensive characterization of guide-specific binding behavior with high sensitivity and reproducibility. We show that TANGO faithfully captures known principles of dCas9 target recognition while revealing previously inaccessible features of guide behavior, including differences in intrinsic affinity, mismatch tolerance, PAM dependence, and RNP assembly. Benchmarking against published and newly generated ChIP-seq datasets demonstrates that while intrinsic DNA-binding specificity is a major determinant of genomic occupancy, chromatin accessibility also influences the intrinsic binding affinity required for dCas9 binding in cells. Finally, we show that TANGO distinguishes working from non-working CRISPRi guides, identifies highly promiscuous guides overlooked by current specificity measures^32–34^, and uncovers unexpectedly strong guide-directed DNA binding in the absence of a PAM. Together, these findings establish diverse aspects of intrinsic DNA recognition as measurable biochemical determinants of dCas9 function, and provide a framework for improving the design of CRISPR technologies that depend on programmable DNA binding.

## RESULTS

### High-throughput array-based assay captures RNP:DNA binding interactions with high sensitivity

To enable precise and high-throughput characterization of dCas9 binding, we developed TANGO (<u>T</u>argeted <u>A</u>rray-based <u>N</u>ucleic acid-<u>G</u>uided <u>O</u>ccupancy), an *in vitro* assay based on the protein-DNA binding microarray (PBM) platform that we and others used extensively to characterize DNA binding by transcription factors and DNA repair proteins^35–41^. Briefly, we use custom-designed DNA libraries synthesized on high-density microarray slides (Agilent) to simultaneously measure dCas9 binding to tens of thousands of 60-bp double-stranded DNA sequences. Each unique sequence is represented in eight or nine replicate spots (depending on the DNA library design; **Methods**), which allows for robust comparisons of dCas9 binding levels among different on-target and off-target DNA sequences. The DNA array is incubated with pre-formed ribonucleoprotein (RNP) complex obtained by incubating purified recombinant dCas9 (C-terminal 6xHis-tagged) with a gRNA of interest (**Figure 1a**). Binding to the DNA sites synthesized on the array is detected with anti-His Alexa488-labelled antibody, with the fluorescence signal at each spot being detected using a high-resolution scanner (**Figure 1b**). Median fluorescence intensities over replicate DNA spots are used to quantify the RNP binding level for each sequence represented in our TANGO DNA library. High-density DNA arrays were chosen as the foundation for our TANGO assay due to their proven ability to generate highly-reproducible DNA binding measurements that quantitatively correlate with the binding affinities of the proteins under investigation, across a wide affinity range^38,39,42–45^.

**Figure 1.**
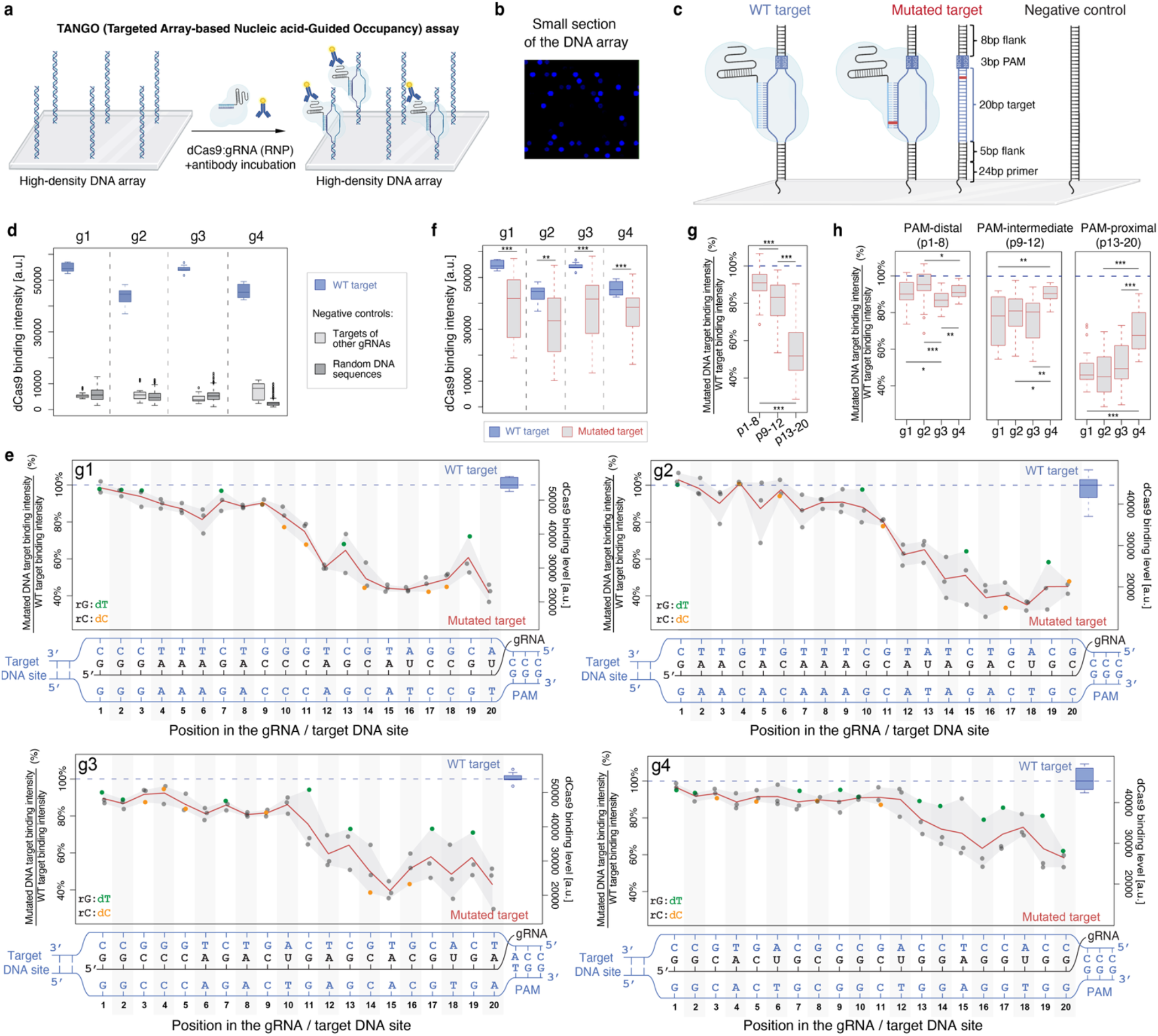
TANGO enables sensitive, high-throughput quantification of dCas9 RNP-DNA binding on high-density DNA arrays. **a**, Overview of the TANGO (Targeted Array-based Nucleic acid-Guided Occupancy) workflow. Pre-formed dCas9:gRNA ribonucleoprotein (RNP) complexes are incubated on high-density double-stranded DNA microarrays, and bound RNP is detected using an anti-His fluorophore-labeled antibody. **b**, Representative image of a small region of the DNA array, after incubation and scanning. Each DNA spot contains millions of identical copies of a particular DNA sequence of interest. The signal at each spot reflects the level of bound dCas9 at that particular DNA sequence, measured using an Alexa488-labeled antibody. **c**, Probe architecture and probe groups printed on TANGO arrays. Each 60-bp probe contains a constant primer region proximal to the array surface and a target region comprising a 20-nt protospacer, a 3-nt PAM, and native genomic flanks. For each gRNA, the TANGO DNA library includes: wild-type (WT) target probes; mutated target probes spanning all single-nucleotide substitutions across the 20-nt protospacer (with PAM and flanks held constant); and negative control (NegCtrl) probes. **d**, gRNA-specific binding to WT target probes (blue), compared to random DNA sites and to targets of other gRNAs (grey); see **Methods** for details. **e**, Mismatch tolerance profiles for the four gRNAs. Left y-axis: percent dCas9 binding for each mutated protospacer position relative to WT (dashed line). Right y-axis: absolute dCas9 binding levels for each mismatch at each position. Points denote individual mismatches, with colored examples highlighting rG:dT (green) and rC:dC (orange). The nucleotide sequences for the target site, the PAM, and the gRNA spacer are shown underneath each profile. The red line shows the average dCas9 binding level at each position (across the three possible mismatches at that position). **f**, Comparison of dCas9 binding to the WT target probe versus the mutated-target probe sets (i.e. probes with 1-nt mutated target site + WT PAM) for each gRNA. **g**, Position dependence of gRNA mismatch tolerance, summarized as percent binding to the mutated-target probes relative to the corresponding WT target (WT set to 100%), grouped into PAM-distal (positions 1-8), PAM-intermediate (positions 9-12), and PAM-proximal (positions 13-20) regions. **h**, gRNA-to-gRNA comparison of mismatch tolerance (computed as percent binding relative to WT) within PAM-distal (left), PAM-intermediate (middle), and PAM-proximal (right) regions, highlighting elevated PAM-proximal permissiveness for g4, i.e. the gRNA with broad off-target binding in cells. Statistical significance according to two-sided Mann-Whitney U-test: *P < 0.05; **P < 0.01; ***P < 0.001.

To benchmark TANGO against existing dCas9 binding data, we first tested four gRNAs (g1, g2, g3, and g4) that were previously characterized using both *in vitro* (i.e. cell-free) and *in vivo* (i.e. cell-based) binding assays^22,28^. ChIP-seq data for the four gRNAs was generated in human embryonic kidney (HEK293T) cells by Kuscu et al.^22^, who identified between 16 and 483 genomic binding locations (i.e. ChIP-seq peaks) per gRNA. For the same four gRNAs, Boyle et al.^30^ used sequencing-coupled filter-binding assays (FBA) to generate *in vitro* binding data for the on-target site as well as all possible 1-nt variants of the protospacer region. We selected these gRNAs because they span a wide range of DNA-binding affinities and mismatch tolerances (according to the *in vitro* FBA data), and a wide range of off-target binding activity (according to the *in vivo* ChIP-seq data), thus providing a well-characterized benchmark for direct comparison to our TANGO measurements.

For each of the four gRNAs, we engineered three groups of DNA probes to be included into our TANGO DNA library and synthesized on the high-density DNA arrays (**Figure 1c**). First, we designed wild-type (WT) target site probes comprising a perfectly matched 20-nt protospacer followed by the native PAM (NGG), embedded in native genomic sequence context (5-nt upstream of the protospacer and 8-nt downstream of the PAM; see **Methods** for details). These probes serve as on-target references (**Figure 1c, left**). Second, for each gRNA of interest we designed “mutated target” probes by mutating each position of the 20-bp DNA target to each of the three alternative base pairs, with the PAM and flanking regions held constant (**Figure 1c, middle**). This comprehensive panel of 3×20 probes allows us to map the positional and nucleotide-specific tolerance of each gRNA to all possible single mismatches in the DNA target, and thus we refer to these probes as the “mismatch tolerance” probe set. Third, we included negative control probes to define a stringent baseline for dCas9 binding at non-specific sites. For this purpose, we generated random DNA sequences with zero, one, two, or three PAMs at different positions within the 60-bp probes (**Methods**; **Table S1**). Finally, we included in the TANGO DNA library a set of randomly selected genomic sites from open chromatin regions in HEK293T cells (according to DNase-seq data from ENCODE, accession number ENCSR000EJR^46^), matching the cell type used by Kuscu et al. for the dCas9 ChIP-seq measurements. All DNA sequences and binding measurements are listed in **Table S2**.

We found that TANGO faithfully recapitulates canonical features of dCas9-DNA recognition, while also revealing intrinsic properties of gRNA g4 that explain its extensive off-target binding in cells, as described in the sections below. Across all four gRNAs tested, perfectly matched (WT) target probes exhibited strong binding, both relative to other gRNA-specific targets present in the TANGO DNA library, and to negative control probes (**Figures 1d, S1a**). TANGO measurements were highly reproducible for all four guides tested, even when assayed at different RNP concentrations (R^2^=0.895-0.959, **Figure S1b**), indicating robust and stable guide-directed dCas9 binding under varied assay conditions. Together, these results demonstrate that TANGO provides a quantitative and reproducible readout of gRNA-specific DNA binding, with minimal binding signal at unrelated sequences, and a clear dynamic range between on-target binding and background.

### TANGO quantifies position- and mismatch-dependent effects of DNA mutations on dCas9 binding

We next quantified how single-nucleotide mismatches between the DNA target and the gRNA spacer influence RNP binding. To do so, we leveraged the “mismatch tolerance” probe sets for each gRNA, encompassing all possible 1-bp substitutions across the 20-bp target site. Each gRNA exhibited a distinct mismatch tolerance profile (**Figure 1e**), with binding effects strongly dependent on both the position and identity of the mismatches. In addition, several consistent patterns emerged. First, for all four gRNAs, mismatched targets showed substantially reduced binding relative to the perfectly matched site (**Figure 1f)**. Second, the magnitude of this reduction varied widely along the protospacer: mismatches in the PAM-proximal region (positions 13-20) caused the strongest decreases in binding, whereas mismatches in the PAM-distal region (positions 1-8) had minimal impacts (**Figure 1e,g**), consistent with prior studies^21,28,30,47–49^. Mismatches in the intermediate region (positions 9-12) showed moderate effects. This partitioning of the 20-bp target sites into PAM-proximal, PAM-intermediate, and PAM-distal regions was guided by prior work^50^, and highlights the dominant contribution of PAM-proximal complementarity to strong dCas9 binding.

Comparing across gRNAs, we found that the relative effects of mismatches (quantified as percent binding relative to the WT target) were broadly similar among gRNAs g1, g2, and g3, with g4 being a clear outlier. Specifically, the mismatch tolerance profiles revealed that g4 is much more promiscuous than the other gRNAs tested, exhibiting much smaller mismatch effects in both the PAM-proximal and PAM-intermediate regions (**Figure 1h**). This elevated mismatch tolerance is consistent with the extensive off-target binding of g4 observed in cells^22^, as described in the sections below. In addition to positional effects, mismatch identity emerged as a key determinant of binding: across all four gRNAs, rG:dT mismatches were the most tolerated, whereas rC:dC mismatches produced the strongest reductions in binding (**Figure 1e**), in agreement with prior studies^31,51–53^.

Collectively, these data demonstrate that TANGO accurately captures the established biochemical determinants of dCas9-DNA recognition. Across multiple gRNAs, the assay reveals high binding specificity, pronounced sensitivity to mismatches in PAM-proximal regions, and clear, quantitative differences across mismatch identities. This strong concordance with prior biochemical understanding provides a robust foundation for subsequent comparisons between TANGO data and dCas9 measurements obtained with alternative assays.

### TANGO captures RNP-DNA binding with higher sensitivity than sequencing-based *in vitro* assays

Few previous studies have characterized the intrinsic DNA-binding specificity of dCas9 RNPs in high throughput^28–31^, with only one study testing gRNAs at scale. In particular, Boyle et al.^28^ used a nitrocellulose filter-binding assay (FBA) coupled to a sequencing-based readout to measure RNP-DNA binding for 90 gRNAs against ∼35,000 on-target and off-target DNA sequences. This dataset includes the four gRNAs (g1–g4) profiled in our initial TANGO experiments (**Figure 1**), along with comprehensive single-mismatch variants of their target sites, enabling a direct side-by-side comparison between FBA and TANGO measurements (**Figure 2a-c**). We found that TANGO and FBA largely agree on the relative binding affinities of dCas9 for the WT target sites of the four gRNAs tested, with g1 and g3 showing higher binding intensities in TANGO, as well as higher DNA fractions bound in FBA, compared to g2 and g4 (horizontal dotted lines in **Figure 2a,b**).

**Figure 2.**
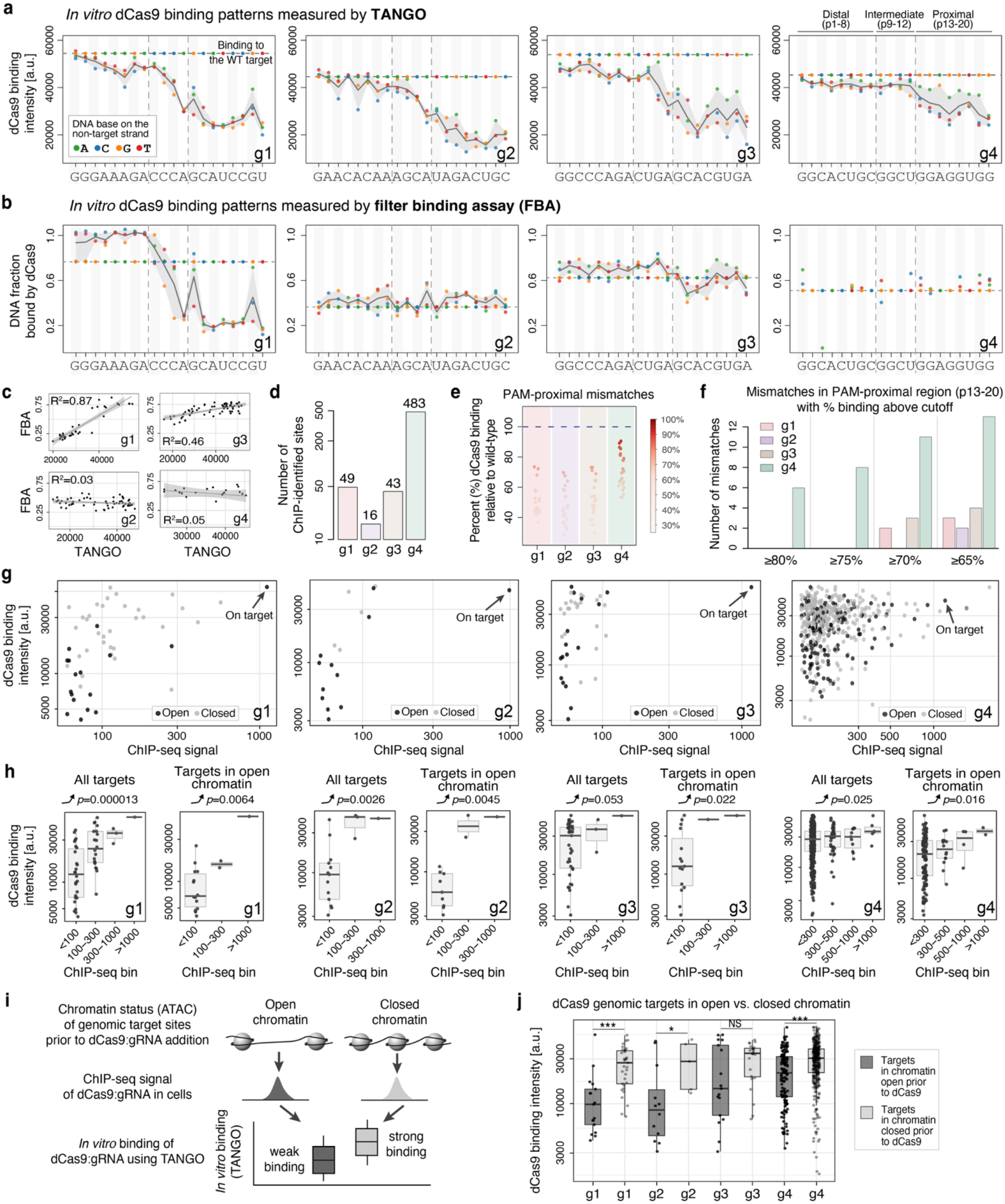
TANGO enables sensitive *in vitro* binding measurements across gRNAs and links intrinsic binding to in-cell occupancy. **a**, *In vitro* mismatch tolerance profiles measured by TANGO for four published guides (g1-g4)^22,28^. Points are individual single-mismatch probes (colored by substituted base on the non-target strand) plotted across target site positions 1-20. Dark grey lines show the mean binding intensity at each position, calculated over the three possible mismatches at that position. Horizontal dotted lines show the binding level at the WT target site. Dashed vertical lines mark PAM-distal (p1-8), PAM-intermediate (p9-12), and PAM-proximal (p13-20) regions. **b**, Similar to panel a, but showing the mismatch profiles derived from the filter binding assay (FBA) data of Boyle et al.^28^. Y-axis shows the fraction of DNA molecules bound, according to the FBA data, for the WT and the mismatched sites. **c**, Correlation between TANGO and FBA measurements across the WT and single-mismatch probes for each gRNA. R²: squared Pearson correlation coefficient. **d**, Number of ChIP-identified dCas9 binding sites (i.e. ChIP-seq peaks) in HEK293T cells for each gRNA. **e,** PAM-proximal mismatch tolerance summarized as percent binding relative to WT. Points are colored by % of WT binding level. **f,** Number of PAM-proximal mismatches retaining binding levels above the indicated thresholds: ≥80%, ≥75%, ≥70%, and ≥65% relative to WT. **g,** Relationship between *in vitro* binding levels measured by TANGO (y-axis) vs. *in vivo* binding signal measured by ChIP-seq (x-axis), for ChIP-identified genomic targets as reported in ^22^. Points are colored by chromatin state (open vs. closed prior to dCas9), and arrows denote the on-target sites. **h**, Aggregate analysis of ChIP-identified targets showing TANGO binding intensity distributions for targets grouped by ChIP-seq signal. Correlation trends are shown for all targets, and for the subset of targets in genomic regions open prior to dCas9 recruitment. P-values according to Jonckheere-Terpstra tests for increasing trend are shown. **i**, Schematic of the chromatin-state stratification analysis. ChIP-identified genomic targets were first classified by pre-existing chromatin accessibility (open vs. closed, based on ATAC-seq data) and then compared by their *in vitro* TANGO binding intensities to assess whether sites bound within closed chromatin show higher intrinsic binding affinity than sites bound in open chromatin. **j**, TANGO binding intensities for ChIP-identified targets stratified by chromatin state (open vs. closed prior to dCas9) for each gRNA. P-values according to one-sided Mann-Whitney U tests are shown: **** p<0.0005, ** p<0.005, * p<0.05.

However, among the four gRNA, FBA only displayed the expected positional mismatch tolerance pattern for g1 (the gRNA with the highest affinity in both assays), characterized by strong reduction in binding for PAM-proximal mismatches and moderate effects in the PAM-intermediate region (**Figure 2a,b**). For this high-affinity gRNA, the TANGO and FBA binding data showed strong correlation across all mismatches in the target region (R^2^=0.87, **Figure 2c**). For the remaining three gRNAs (g2, g3, and g4), TANGO robustly resolved the RNP’s tolerance to mismatches across the target site (**Figure 2a**), whereas the corresponding FBA-derived mismatch tolerance profiles were largely flat (**Figure 2b**), suggesting little to no apparent specificity according to FBA. Furthermore, in the case of g4, FBA binding measurements were missing for many of the mismatched targets, as they fell below the assay’s detection limit^28^. This lower or undetectable level of binding in the FBA assay is inconsistent with ChIP-seq data^22^, which we describe in the next section.

Overall, this side-by-side comparison demonstrates that compared to FBA, which is the only existing *in vitro* technology that has been used at scale for dCas9 binding measurements, TANGO more sensitively and faithfully captures RNP-DNA binding across gRNAs spanning a range of affinities and specificities. This enhanced sensitivity likely reflects key features of the TANGO array-based platform, including (i) a direct binding readout that does not rely on target capture or sequence amplification, (ii) multiple technical replicates per probe that reduce spot-level noise, and (iii) a broad dynamic range that enables detection of weaker binding interactions.

### Intrinsic DNA-binding specificity captured by TANGO reflects dCas9 occupancy in cells

Having established that TANGO sensitively captures the intrinsic DNA-binding specificity of dCas9 across diverse gRNAs, we next tested whether these measurements reflect dCas9 binding in cells, as determined by ChIP-seq assays. Genome-wide dCas9 ChIP-seq data in HEK293T cells is available for the four gRNAs described above^22^. The number of ChIP-seq peaks identified across the genome ranges from 16 (for g2) to 483 (for g4), indicating substantial differences in off-target binding (**Figure 2d**).

Across all four gRNAs, we found good agreement between *in vitro* binding as captured by our array-based TANGO assay and *in vivo* binding as measured by ChIP-seq (**Figure 2g,h**), consistent with prior findings for transcription factor proteins profiled using a similar array-based platform^39^. As expected given the complexity of the nuclear environment relative to a cell-free system, site-by-site correlations between TANGO and ChIP-seq are modest (**Figure 2g**). However, aggregate quantile analyses of the dCas9 ChIP-seq data displayed significant correlations with our TANGO measurements (**Figure 2h**). Importantly, TANGO correctly ranked the on-target sites (**Figure 2g**, black arrows) among the highest-affinity binding sites for all gRNAs. As expected, we also observed instances where off-target sites exhibited strong *in vitro* binding but weaker ChIP-seq signal (**Figure 2g**, upper-left points). This is largely attributable to limited chromatin accessibility in cells, as many of these sites reside in regions that are closed prior to dCas9 recruitment (light grey points in **Figure 2g**).

Notably, we did not observe any genomic targets with strong ChIP-seq signal but weak *in vitro* binding (**Figure 2g**), indicating that high intrinsic binding affinity, as captured by TANGO, is a prerequisite for robust occupancy in cells. Consistent with this, targets detected in both ChIP-seq replicates exhibited higher TANGO binding intensities than those identified in only a single replicate (**Figure S2g,h**), suggesting that intrinsic binding strength contributes to more reproducible and robust genomic binding in cells. Together, these results support the future use of TANGO-derived metrics to guide optimal gRNA selection, enabling the systematic de-prioritization of guides with low affinity and/or specificity in large-scale epigenetic screens.

An additional pattern emerged upon stratifying genomic target sites by chromatin accessibility prior to dCas9 recruitment (open versus closed; **Figure 2i**). Interestingly, off-target sites located in closed chromatin (prior to dCas9) exhibited significantly higher *in vitro* binding levels compared to off-target sites located in open chromatin (**Figure 2j**). This relationship persisted after normalizing TANGO binding intensities to ChIP-seq signal (**Figure S2e,f)**, indicating that this finding is not simply driven by differences in ChIP enrichment. Together, these results suggest a model in which RNPs can readily engage lower-affinity sites in accessible chromatin, but binding within closed, nucleosome-occupied regions requires substantially higher intrinsic affinity to overcome chromatin barriers. This behavior parallels that observed for pioneer transcription factors, which bind a broad range of sites in open chromatin, but engage closed chromatin and initiate remodeling predominantly at higher-affinity binding sites^54,55^.

### Mismatch tolerance measured by TANGO identifies gRNA with broad off-target binding in the cell

We next asked whether mismatch tolerance profiles measured by TANGO could identify gRNAs prone to widespread off-target binding in cells. To address this, we focused on mismatches within the PAM-proximal region (bases 13-20 in the protospacer) of the target sites for the four gRNAs described above. For each mismatch, we calculated dCas9 binding to the mismatched sequence relative to the perfectly matched target and compared these values across gRNAs (**Figure 2e**). We then quantified the number of PAM-proximal mismatches exceeding defined binding thresholds (65%, 70%, 75%, 80%) relative to the wild-type site (**Figure 2f**), providing summary metrics of mismatch tolerance for each gRNA. These analyses showed that g1, g2, and g3 exhibit strong discrimination against PAM-proximal mismatches, with all such mismatches reducing binding to ≤75% of the WT target, and only 2-4 PAM-proximal mismatches retaining ≥65% of WT binding (**Figure 2f**). This pronounced intolerance, which indicates high intrinsic DNA-binding specificity, is in great agreement with the relatively small number of off-target sites identified by ChIP-seq for these gRNAs (**Figure 2d**). In contrast, gRNA g4 maintains high levels of binding across many PAM-proximal mismatches (**Figure 2e,f**), with 14 mismatches retaining ≥65% of WT binding (**Figure 2f**). This high tolerance to mismatches is fully consistent with the extensive off-target binding observed for g4 in cells (483 ChIP-seq peaks, **Figure 2d**), indicating that TANGO can effectively flag promiscuous gRNAs with high off-target potential. These trends remained robust when expanding the analysis to include PAM-intermediate positions, and when using absolute binding thresholds instead of percent-of-WT thresholds (**Figure S2b-d**), confirming that g4’s elevated permissiveness relative to the other gRNAs is not dependent on a specific metric. Together, these results demonstrate that the intrinsically higher mismatch tolerance of g4 relative to g1-g3, as measured by TANGO (**Figure 2e,f)**, directly corresponds to its markedly increased off-target binding in cells, where g4 exhibited an order of magnitude more off-target sites than the other gRNAs.

Focusing on genomic off-target sites identified by ChIP-seq for g1-g4, we next asked whether these sites preferentially contain mismatches that are well tolerated according to our *in vitro* TANGO measurements. Notably, genomic off-targets with only a single mismatch relative to the on-target sequence are expected to be very rare, as the human genome contains only a small fraction (at most ∼0.37%, if all genomic 20-mers were unique) of all possible 20-mers (the genome can accommodate at most ∼3x10^9^ of the possible 4^20^ or ∼1.1x10^12^ 20-mers). Accordingly, most genomic off-targets contain multiple mismatches relative to the on-target site. The combinatorial effects of these mismatches are not fully understood and are not simply additive, i.e., the total binding penalty at an off-target site does not equal the sum of penalties from individual mismatches. This was both observed in our TANGO data (**Figure S2i**) and reported previously^56^. Close analysis of the ChIP-seq data revealed that ChIP-identified off-target sites (**Figure S3**) generally contain relatively few mismatches in the PAM-proximal and PAM-intermediate regions, with the notable exception of g4, which also exhibited high intrinsic tolerance to such mismatches in TANGO. When mismatches were present in these critical regions, they tended to correspond to mismatch types that retained stronger binding in TANGO (**Figure S3**), indicating that the intrinsic mismatch tolerance of gRNAs is a key quantitative determinant of which off-target sites are ultimately occupied by dCas9 in cells.

More broadly, these findings highlight how the intrinsic tolerance/intolerance of gRNAs, as captured by TANGO, could be used in future studies to systematically flag and exclude gRNAs with high off-target binding potential from CRISPRi/a studies. We note, however, that single-mismatch tolerance profiles are not sufficient to predict dCas9 binding to genomic off-targets, even in cell-free conditions (**Figure S2i**), and more TANGO data is needed to learn predictive models of dCas9 off-target binding both in cell-free and cellular systems.

### TANGO validates on-target binding for active CRISPRa guides and flags mismatch-permissive outliers confirmed by ChIP-seq

Strong and specific dCas9 binding to its genomic target sites is a prerequisite for CRISPRa and CRISPRi activity. To test whether TANGO can robustly capture the binding specificity of guides with demonstrated activity in cells, we profiled eleven CRISPRa gRNAs that activate the *FOXO4* gene in primary astrocytes^57^ (**Figure 3a,b;S4a**). In TANGO assays, all eleven gRNAs exhibited strong and specific binding to their on-target sequences (**Figure 3c, S4b**) and displayed the expected positional mismatch tolerance profiles, characterized by strong reductions in binding for PAM-proximal mismatches (positions 13-20), moderate effects in the PAM-intermediate region (positions 9-12), and minimal effects in the PAM-distal region (positions 1-8) (**Figure 3d-h, S4c**). These findings demonstrate that gRNAs capable of driving robust CRISPRa activity in cells also exhibit strong and sequence-specific intrinsic binding in TANGO, consistent with our observations from the ChIP-benchmarking gRNAs (g1-g4) that sequence-intrinsic RNP binding is a prerequisite for in-cell occupancy and, ultimately, for regulatory activity.

**Figure 3.**
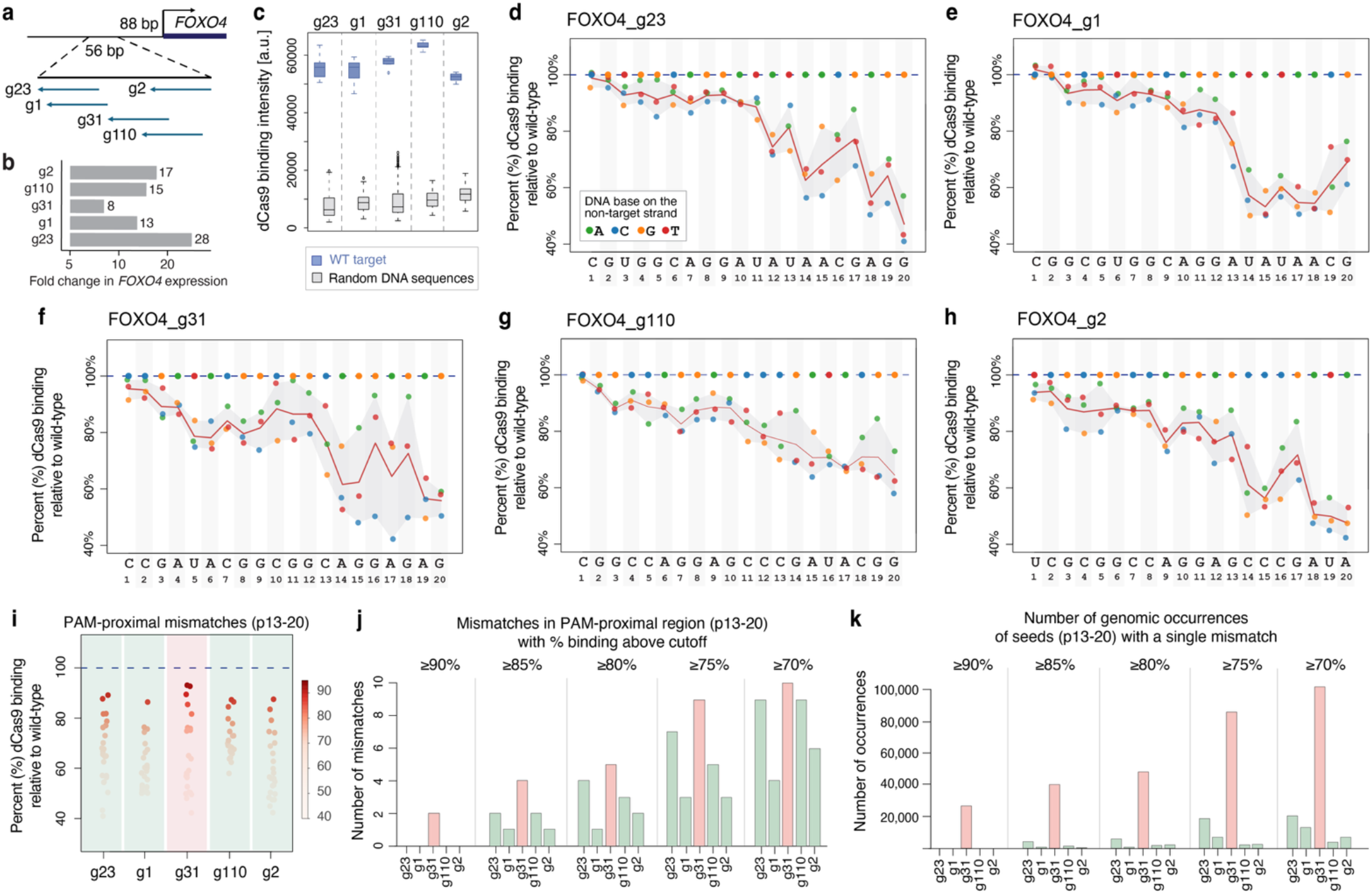
TANGO validates on-target binding for active *FOXO4* CRISPRa guides and identifies a mismatch-permissive outlier. **a**, Five CRISPRa gRNAs (g23, g1, g31, g110, g2) targeting a contiguous 56-bp region in the *FOXO4* promoter. **b,** CRISPRa activity of the five gRNAs in primary astrocytes^58^, shown as fold change in *FOXO4* expression. **c**, TANGO data showing gRNA-specific binding to WT target probes (blue) compared to random DNA sequences (grey). **d-h**, Mismatch tolerance profiles for the five gRNAs, plotted as percent dCas9 binding for each mutated protospacer position relative to WT (dashed line). Points denote individual mismatches (colored by substituted base) and the line shows the average dCas9 binding level at each position (across the three possible mismatches at that position). **i**, PAM-proximal mismatch tolerance summarized as percent binding relative to WT. Points are colored by % of WT binding level. The gRNA with the highest mismatch tolerance, g31, is highlighted using a light red background. **j,** Number of PAM-proximal mismatches retaining binding levels above the indicated thresholds: ≥90%, ≥85%, ≥80%, ≥75%, and ≥70% relative to WT. **k,** Number of occurrences across the human genome (hg38) of 8-mer seeds (positions 13-20) with a single mismatch and that retained binding levels above the indicated thresholds relative to WT. Guide g31 stands out as a promiscuous gRNA with high potential for widespread binding across the genome.

To further explore the relationship between intrinsic binding specificity and regulatory activity, we focused on five gRNAs targeting a contiguous 56-bp region of the *FOXO4* promoter that produced gene activation levels ranging from 8-fold to 28-fold^57,58^ (**Figure 3a,b**). These guides were originally tested in a large-scale CRISPRa screen of human transcription factor genes aimed at discovering factors capable of reprogramming astrocytes toward neuron-like states^57^. Although all five guides exhibited strong and specific on-target binding in TANGO (**Figure 3c**), the mismatch tolerance profile of g31 emerged as a clear outlier (**Figure 3f**). In particular, g31 retained relatively high binding across a larger set of PAM-proximal mismatches compared to the other guides (**Figure 3i,j**), indicating elevated intrinsic permissiveness toward potential off-target sites. In addition, scanning the genome for strongly-bound single-mismatch seed sequences (i.e. 8-mers that are only one nucleotide different from the PAM-proximal region of the on-target site, followed by an NGG PAM), we found orders of magnitude more genomic matches for g31 compared to the other four gRNAs (**Figure 3k**). We do not expect all these genomic sites to be bound by the g31 RNP in cells, as many of them may have highly detrimental mismatches in the PAM-intermediate and PAM-distal regions, or may not be strong enough for dCas9 to overcome nucleosome occupancy. Nevertheless, our results suggest that g31 has a high potential for widespread off-target binding across the genome. Interestingly, g31 also produced the weakest transcriptional activation in the CRISPRa screen compared to the other promoter gRNAs (**Figure 3b**), consistent with the possibility that extensive off-target binding of g31 reduces effective occupancy and activation level at the intended on-target site in the *FOXO4* promoter.

To test the off-target binding activity of g31 in cells, we performed dCas9 ChIP-seq in astrocytes transduced with lentivirus expressing g31 together with FLAG-tagged ^VP64^dSpCas9^VP64^, as described previously^58^ (**Methods**). Using a stringent peak-calling threshold (q-value < 10e-5), we identified 44,732 ChIP-seq peaks that were reproducible across three biological replicates (**Methods**; **Table S3**). This is a surprisingly large number of peaks given that g31 is part of the Calabrese gRNA library recommended for CRISPRa screens^33^, and that g31’s predicted specificity scores from GuideScan2^32^ and from IDT’s CRISPR-Cas9 guide RNA design tool^34^ do not suggest an unusually promiscuous guide. Instead, these scores are comparable to those of the other guides targeting the same genomic region (**Table S1**).

As expected, the intended g31 target site in the *FOXO4* promoter showed strong ChIP-seq signal, quantified as the read pileup within ±150-bp of the peak summit (**Figure 4a**). However, 34 off-target sites showed an even higher ChIP-seq read pileup than the on-target site (**Figure S5; Table S3**), with all of these high-occupancy off-target peaks containing either the perfect g31 seed sequence (CAGGAGAG) or a 1-mismatch seed, immediately followed by an NGG PAM. Across the full set of 44,732 peaks, 10.6% contained the perfect seed sequence, and 84.6% contained a seed with 0-2 mismatches relative to the gRNA (**Figure 4b**). We next examined whether chromatin accessibility influences the sequence requirements for off-target binding. Using ATAC-seq data from astrocyte cells without g31 present^57^, we found that ChIP-seq peaks located in closed chromatin had seed sequences with fewer mismatches than peaks in accessible chromatin (**Figure 4b**). This observation is consistent with our analysis of the g1-g4 ChIP-benchmarking guides, which showed that off-target sites in closed chromatin require higher intrinsic binding affinity than those in open chromatin (**Figure 2j**).

**Figure 4.**
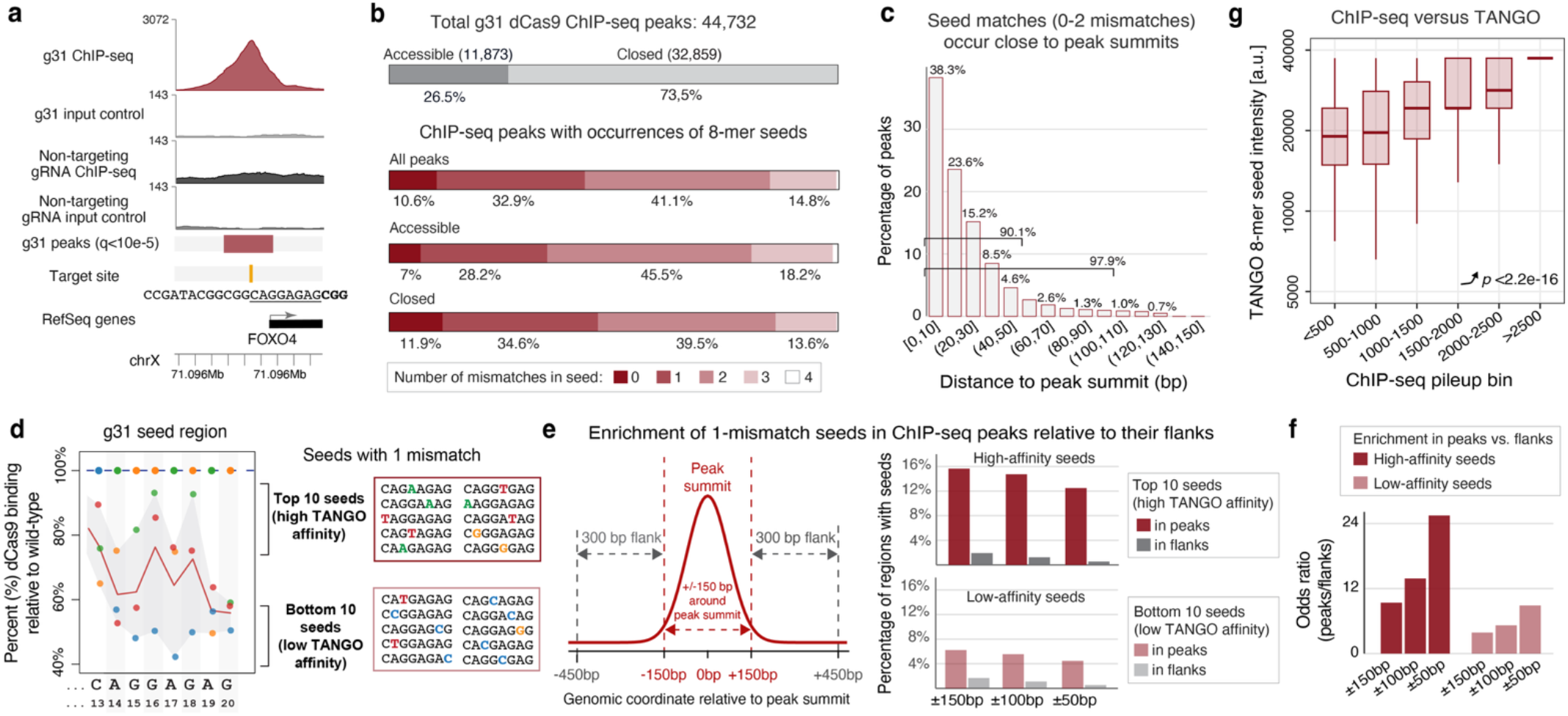
ChIP-seq data confirms widespread off-target binding by promiscuous gRNA flagged by TANGO. **a**, ChIP-seq data, at the on-target site in the *FOXO4* promoter, for the g31 guide. **b**, Number of ChIP-seq peaks identified for dCas9+g31 in astrocytes (**Methods**). **c**, Occurrences of 8-mer seed sequences with 0-2 mismatches within ChIP-seq peaks. Seed matches are located in close proximity to the peak summits. X-axis intervals show the range of distances to the peak summit for each bin. **d**, Single-mismatch seeds, i.e. 8-mers that are 1-nt different from the 8-bp PAM-proximal region of the perfect g31 target site, were grouped based on their TANGO binding levels. We focused on the top ten (highest-affinity) and the bottom ten (lowest-affinity) 1-mismatch seeds, as shown. **e**, Left: enrichment of 8-mer seeds was computed in the ChIP-seq peaks vs. their flanking regions, as shown. Right: percentage of peaks (red) or flanking regions (grey) with occurrences of high-affinity seeds (top) vs. low-affinity seeds (bottom). The percentages were computed relative to the full set of 44,732 peaks. Enrichments were assessed in the regions ±150-bp, ±100-bp, and ±50-bp around the peak summits or the centers of the flanking regions. **f**, Enrichment values, shown as odd-ratios of occurrences in the peaks vs. the flanks, for the high-affinity and low-affinity seeds. **g**, Relationship between *in vitro* binding levels measured by TANGO (y-axis) vs. *in vivo* binding signals measured by ChIP-seq (x-axis). Only ChIP-seq peaks that overlap ATAC-seq peaks in astrocytes, and containing a single occurrence of an 8-mer seed with 0-2 mismatches were considered (**Methods**). Y-axis values represent TANGO binding intensities for the 0-1 mismatch seeds, and predicted binding intensities for 2-mismatch seeds, assuming additive penalties for the two mismatches. P-value shown is according to a Jonckheere-Terpstra test for increasing trend.

The high quality of the g31 ChIP-seq data was further supported by the fact that most seed occurrences were positioned very close to the peak summits. For example, 90.1% of peaks have a seed (with 0-2 mismatches) immediately followed by NGG, within 50-bp of the summit, while 97.9% have such a seed within 100-bp of the summit (**Figure 4c**). These percentages were even higher for peaks containing perfect seed matches, as expected (**Figure S4e**), and remained largely unchanged when peak calling was repeated using a more permissive q-value threshold of 0.001 (**Figure S4f,g**).

Having established the high quality of the g31 ChIP-seq data, we next asked whether seed sequences with high affinity according to our TANGO data were more enriched in the g31 ChIP-seq peaks compared to low-affinity seed sequences. To address this question, we grouped 1-mismatch seeds into different categories based on their TANGO binding intensities. First, we focused on the ten highest-affinity and the ten lowest-affinity 1-mismatch seeds (**Figure 4d**), and searched for their occurrences within ChIP-seq peaks and their flanking regions (**Figure 4e**, left), focusing on the 32.9% of peaks that lacked a perfect seed match but contained seeds with a single mismatch (**Figure 4b**). We found that a higher fraction of peaks (∼16%) contained high-affinity 1-mismatch seeds compared to low-affinity 1-mismatch seeds (∼6% of peaks), as shown in **Figure 4e** (dark red vs. light red bars). In contrast, the fractions of flanking control regions with high-affinity vs. low-affinity seeds were uniformly low and not significantly different between the two groups of seeds (**Figure 4e**, grey bars). We note that both sets of seeds were significantly enriched within ChIP-seq peaks relative to the flanking regions (Fisher’s exact test p-value < 10^-200^, **Table S4**). However, the enrichment odds ratio was much larger for high-affinity than low-affinity seeds, particularly within ±50 bp of the peak summit (**Figure 4f**). The same trend was observed when the 1-mismatch seeds were partitioned into high-, medium-, and low-affinity groups based on their TANGO binding intensities (**Figure S4h**). High-affinity seeds showed the strongest enrichment in ChIP-seq peaks, followed by medium- and then low-affinity seeds (**Figure S4i**).

Finally, we asked whether the overall agreement between the *in vitro* binding captured by TANGO and the *in vivo* binding measured by ChIP-seq, as we observed for the g1-g4 ChIP-benchmarking gRNAs (**Figure 2**), also extends to the large set of off-target sites observed for the highly promiscuous g31 gRNA. Because TANGO measurements are not available for the tens of thousands of genomic off-target sites bound by g31 in cells, we instead used as proxies the measured or estimated binding intensities of the corresponding 8-mer seed sequences. For seeds with 0 or 1 mismatches we used direct TANGO measurements, while for seeds with 2 mismatches we estimated TANGO binding intensity by assuming additive mismatch penalties (**Methods**). We focused on the 25,148 ChIP-seq peaks containing a single occurrence of a 0-2 mismatch seed followed by an NGG PAM within ±100 bp of the peak summit (**Table S3)**. Similar to the g1-g4 benchmarking guides (**Figure 2h**), ChIP-seq peaks were grouped according to their read pileup, and the distribution of TANGO seed binding intensities was examined across the ChIP-seq bins.

We observed a clear agreement between g31’s intrinsic binding specificity captured by TANGO and its genome-wide occupancy measured by ChIP-seq (**Figure 4g, Figure S4j**), with this relationship being most significant for sites located in chromatin that was accessible prior to dCas9 recruitment (Jonckheere-Terpstra test for increasing trend, p<2.2e-16, **Figure 4g**). Consistent with our analyses of the g1-g4 ChIP-benchmarking guides (**Figure 2g,j**), we did not observe genomic sites with strong ChIP-seq signal but weak intrinsic binding (**Figure 4g**, **S4j**; **Table S3**), further supporting the conclusion that high intrinsic binding affinity is a requirement for robust occupancy in cells. We also found that off-target sites located in closed chromatin prior to dCas9 recruitment exhibited higher intrinsic binding affinities than sites in accessible chromatin (one-sided Mann-Whitney U test p=2.7e-67, **Figure S4k**), consistent with the notion that stronger RNP-DNA interactions are required to overcome chromatin barriers.

Together, our results on the g31 gRNA show that the intrinsic DNA-binding specificity captured by TANGO is reflected in the genome-wide occupancy of such a highly promiscuous guide. More broadly, the results demonstrate that TANGO can identify guides with a high propensity for widespread off-target binding, and that intrinsic DNA-binding specificity is a major determinant of genome-wide dCas9 occupancy in cells.

### TANGO distinguishes working from non-working CRISPRi guides based on their intrinsic DNA-binding properties

Having shown that TANGO robustly captures the DNA-binding specificity of gRNAs with demonstrated regulatory activity in cells, and correctly identifies highly promiscuous guides like g31, we next asked whether TANGO could distinguish active from inactive CRISPRi guides and provide insights into why some gRNAs fail to modulate gene expression even when neighboring gRNAs targeting the same regulatory element are highly active. This question is particularly relevant for CRISPRi/a screens that target distal regulatory elements, for which we and others have consistently found that only about 10-20% of gRNAs show activity on target gene expression^7,8,59–62^, even when focusing on regulatory elements where at least one gRNA showed activity.

To address this question, we analyzed two groups of gRNAs previously tested in CRISPRi screens of distal regulatory elements. The first group consists of four gRNAs (HS3.g3, HS3.g7, HS3.g27, and HS3.g33) targeting a 46-bp window within the HS3 DNase I hypersensitive site that regulates *HBE1*^59^ (**Figure 5a**). The second group consists of four gRNAs (chr11.g2, chr11.g7, chr11.g8, and chr11.g10) targeting a 100-bp DNase I hypersensitive region (chr11.1734) that regulates *LMO2*^63^ (**Figure 5b**). Within each group, only two of the four gRNAs produced a significant reduction in target gene expression (p<0.05), whereas the remaining two had no detectable effect (**Figure 5c**). We therefore refer to these guides as “working” (labeled in green) and “non-working” (labeled in red), respectively.

**Figure 5.**
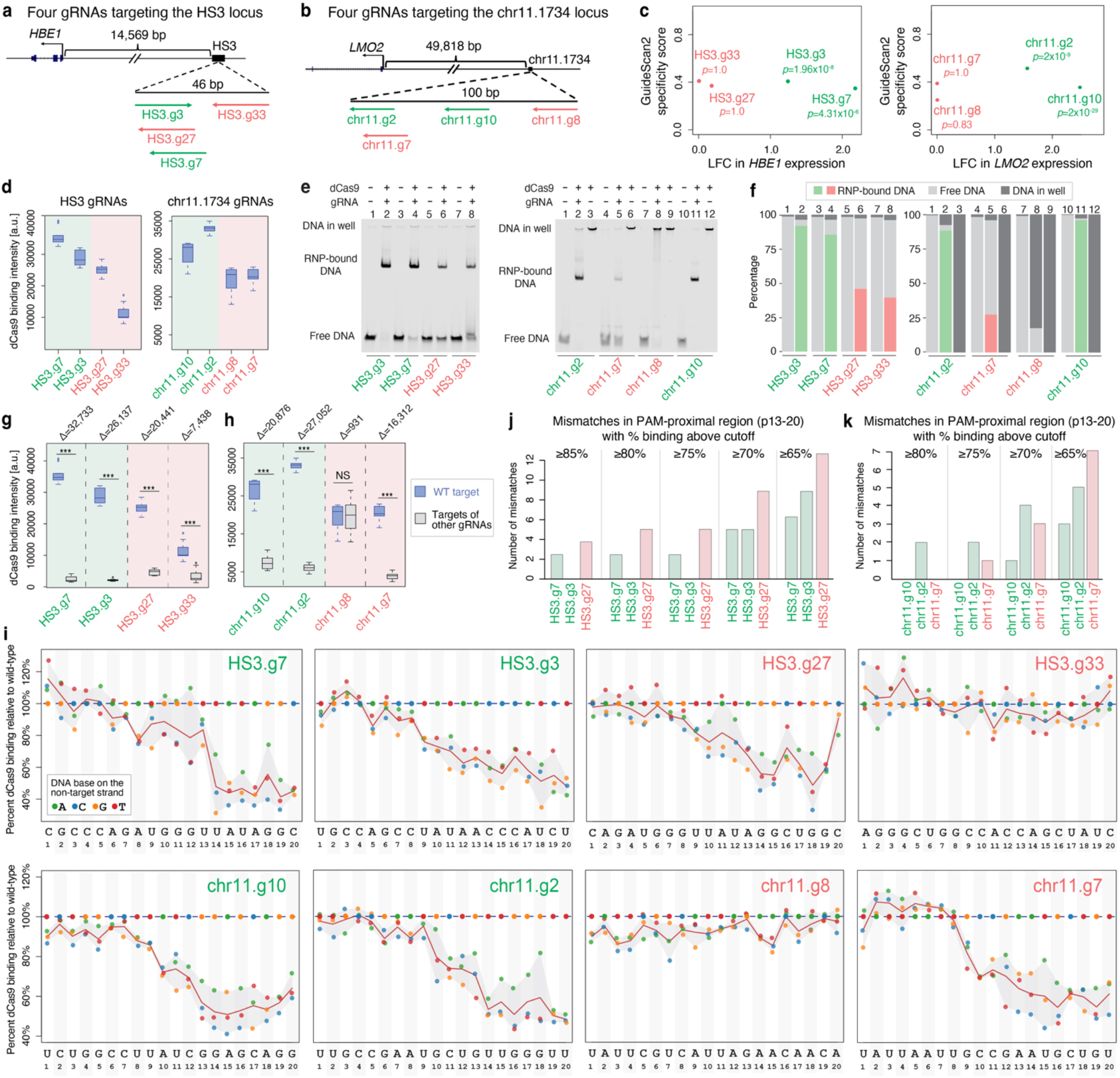
TANGO distinguishes working from non-working CRISPRi guides at two cis-regulatory loci. **a**, Schematic of the HS3 regulatory region of the *HBE1* gene, and the four CRISPRi gRNAs tested at this locus. Working gRNAs, which significantly repressed *HBE1* in the CRISPRi screen^59^, are shown in green; non-working gRNAs are shown in red. Arrows show the genomic orientation of the gRNAs. **b,** Similar to panel a, but for the chr11.1734 regulatory region of the *LMO2* gene^63^. **c,** GuideScan2 specificity scores of the tested gRNAs, plotted against log fold change (LFC) in *HBE1* (left) or *LMO2* (right) expression in the CRISPRi screens. P-values from the CRISPRi screens^59,63^ are shown. **d**, TANGO binding intensities for the HS3 and chr11.1734 gRNAs measured at their on-target probes (**Table S2**). **e**, EMSA gels showing binding of the eight gRNAs to their on-target DNA sites (**Methods**). The EMSA for the chr11.1734 gRNAs includes lanes where only the dCas9 protein was added to the target DNA sites, without any gRNAs. **f,** Quantification of the EMSA results in panel e, using ImageJ (**Methods**). **g-h,** TANGO binding intensities for the HS3 and chr11.1734 gRNAs measured at their on-target probes (blue) versus control probes (here, the targets of other, non-overlapping gRNAs printed on the same array; gray) (**Table S2**). Differences (Δ) are shown between median TANGO binding levels at the on-target vs. the control sites. ***p<0.0005 according to a two-sided Mann-Whitney U test. NS = non-significant. **i**, TANGO mismatch tolerance profiles for the eight gRNAs. Points are individual single-mismatch probes (colored by substituted base on the non-target strand) plotted across target site positions 1-20. Red lines show the mean binding intensity at each position, calculated over the three possible mismatches at that position. Horizontal dotted lines show the binding level at the WT target site**. j-k,** PAM-proximal mismatch tolerance data summarized as the number of PAM-proximal single mismatches (positions 13-20) that retained binding above the indicated thresholds (≥65%, ≥70%, ≥75%, ≥80%, ≥85%) relative to the wild-type site.

Importantly, within each group, the working and non-working guides have comparable predicted specificity scores (**Figure 5c; Table S1**), and they target nearby sites within the same accessible regulatory element, with similar orientation and distance relative to the target gene’s transcription start site. Thus, the marked differences in CRISPRi activity cannot be readily explained by obvious differences in guide specificity scores or epigenetic context. Although local cellular features, such as transcription factor occupancy, DNA methylation, or differences in guide expression, may also affect guide activity in the cell, here we asked whether the two classes of guides (working vs. non-working) differ in their intrinsic DNA-binding properties as measured by TANGO.

In both gRNA groups, TANGO distinguished working from non-working gRNAs based on their intrinsic DNA-binding affinities and/or specificities. In particular, the working gRNAs (HS3.g7, HS3.g3, chr11.g10, and chr11.g2) exhibited higher binding affinities for their intended target sites than the corresponding non-working gRNAs (**Figure 5d**, two-sided Mann-Whitney U test p=2.138e-08 for the HS3 guides and 1.497e-07 for the chr11.1734 guides). To independently validate the on-target binding strengths for the eight gRNAs using an orthogonal approach, we performed electrophoretic mobility shift assays (EMSAs) using all eight guides and their corresponding wild-type target sequences. We used 60-bp fluorescently-labeled DNA probes carrying the WT target sequence for each of the eight gRNAs, matching the sequence context used in TANGO. Consistent with the TANGO results, the working gRNAs showed stronger DNA binding than the non-working gRNAs, with chr11.g8 showing no detectable DNA binding (**Figure 5e,f**).

Second, we found that the working gRNAs had higher DNA-binding specificities than the non-working gRNAs, as they exhibited larger differences in TANGO binding levels between the on-target and the control sites (**Figure 5g,h**). Furthermore, TANGO identified one of the non-working guides, chr11.g8, as a clear outlier that showed essentially no guide-specific DNA binding (**Figure 5h**). This observation raised the possibility that chr11.g8 fails to assemble efficiently with dCas9, such that the binding detected by TANGO is attributable primarily to the intrinsic DNA-binding activity of dCas9 rather than guide-directed recognition. This hypothesis was confirmed in our gel-shift assay, where binding of dCas9 to the chr11.g8 target sequence was indistinguishable in the presence vs. the absence of the chr11.g8 gRNA, with most of the fluorescently-labeled chr11.g8 DNA sample retained near the well of the gel (**Figure 5e,f**). Thus, EMSA provides independent validation of TANGO, while also supporting that chr11.g8’s lack of guide-specific binding signal in TANGO is likely a result of impaired RNP formation.

To further investigate differences between working and non-working gRNAs, we next compared their TANGO-derived mismatch tolerance profiles (**Figure 5i**). We first examined mismatch-dependent binding across the full protospacer to determine whether each gRNA exhibited a clear guide-specific mismatch signature, characterized by progressively reduced binding upon introduction of mismatches, particularly in the PAM-proximal region (positions 13-20). Two of the non-working gRNAs, HS3.g33 and chr11.g8, displayed essentially flat mismatch tolerance profiles, including across the PAM-proximal region, indicating little or no guide-directed DNA recognition. For chr11.g8, this observation is consistent with the lack of guide-specific on-target DNA binding detected by TANGO (**Figure 5h**) and confirmed by EMSA (**Figure 5e,f**), which together indicate inefficient dCas9 RNP assembly. In contrast, although HS3.g33 binds its intended target modestly better than the control sites (**Figure 5g**) and does not show evidence of defective RNP assembly **(Figure 5e,f**), it likewise lacks a discernible guide-specific mismatch tolerance signature, suggesting that it fails to direct efficient sequence-specific DNA recognition through a different mechanism.

The remaining two non-working gRNAs, HS3.g27 and chr11.g7, showed the expected guide-specific mismatch profiles but displayed substantially greater tolerance to PAM-proximal mismatches than the corresponding working gRNAs. Specifically, they retained high binding across a larger number of PAM-proximal mismatches (**Figure 5j,k**), and these strongly bound mismatch seeds occurred more frequently throughout the human genome than the strongly bound mismatched seeds of the working guides (**Figure S6b,c**), suggesting a greater potential for widespread off-target binding. Together with their reduced on-target binding affinities (**Figure 5d**), these observations suggest that the lack of CRISPRi activity of HS3.g27 and chr11.g7 reflects a combination of weaker on-target binding and more permissive sequence recognition, increasing the likelihood of competing off-target interactions in cells.

Together, these results demonstrate that TANGO distinguishes working from non-working CRISPRi gRNAs based on their intrinsic DNA-binding affinity and specificity, identifies gRNAs with inefficient dCas9 RNP assembly, and provides mechanistic insights into why individual gRNAs fail despite targeting regulatory elements that are successfully perturbed by neighboring guides. As TANGO measurements become available for larger numbers of gRNAs, these data should enable the identification of general DNA-binding features associated with poor CRISPRi performance, facilitating the exclusion of suboptimal gRNAs during CRISPRi library design, and the selection of more effective candidates for epigenome editing applications.

### PAM-independent binding reveals an additional layer of guide permissiveness

Next, we used TANGO to ask whether guide behavior is influenced not only by mismatch tolerance within the protospacer but also by sequence variation within the PAM region. To address this question, we included in our TANGO DNA libraries for the eight CRISPRi gRNAs (**Figure 5a,b**) DNA probes that maintained the wild-type 20-bp target sequence while varying the adjacent PAM sequence (**Figure 6a**). For each gRNA, we compared binding to the native NGG PAM, the three alternative NGG PAM variants, and a set of “no-PAM” sequences (**Figure 6b**; **Methods**).The no-PAM probes included sequences where the NGG PAM was replaced by a random trinucleotide (5’-TAC-3’), as well as sequences where the first PAM nucleotide was identical to that of the corresponding wild-type PAM but the downstream GG dinucleotide was replaced with alternative dinucleotides, excluding the commonly reported non-canonical PAMs NAG and NGA^64–66^. Across all eight gRNAs, the native PAM and the alternative NGG PAM variants exhibited similar binding affinities (**Figure 6b**, dark blue vs. light blue boxes). Interestingly, for most guides the native PAM was not the highest-affinity NGG variant (**Table S2**), indicating that nucleotide identity at the first base of the PAM can measurably influence dCas9 binding despite all four NGG PAMs (AGG, CGG, GGG, and TGG) supporting efficient target recognition^48,64,67^.

**Figure 6.**
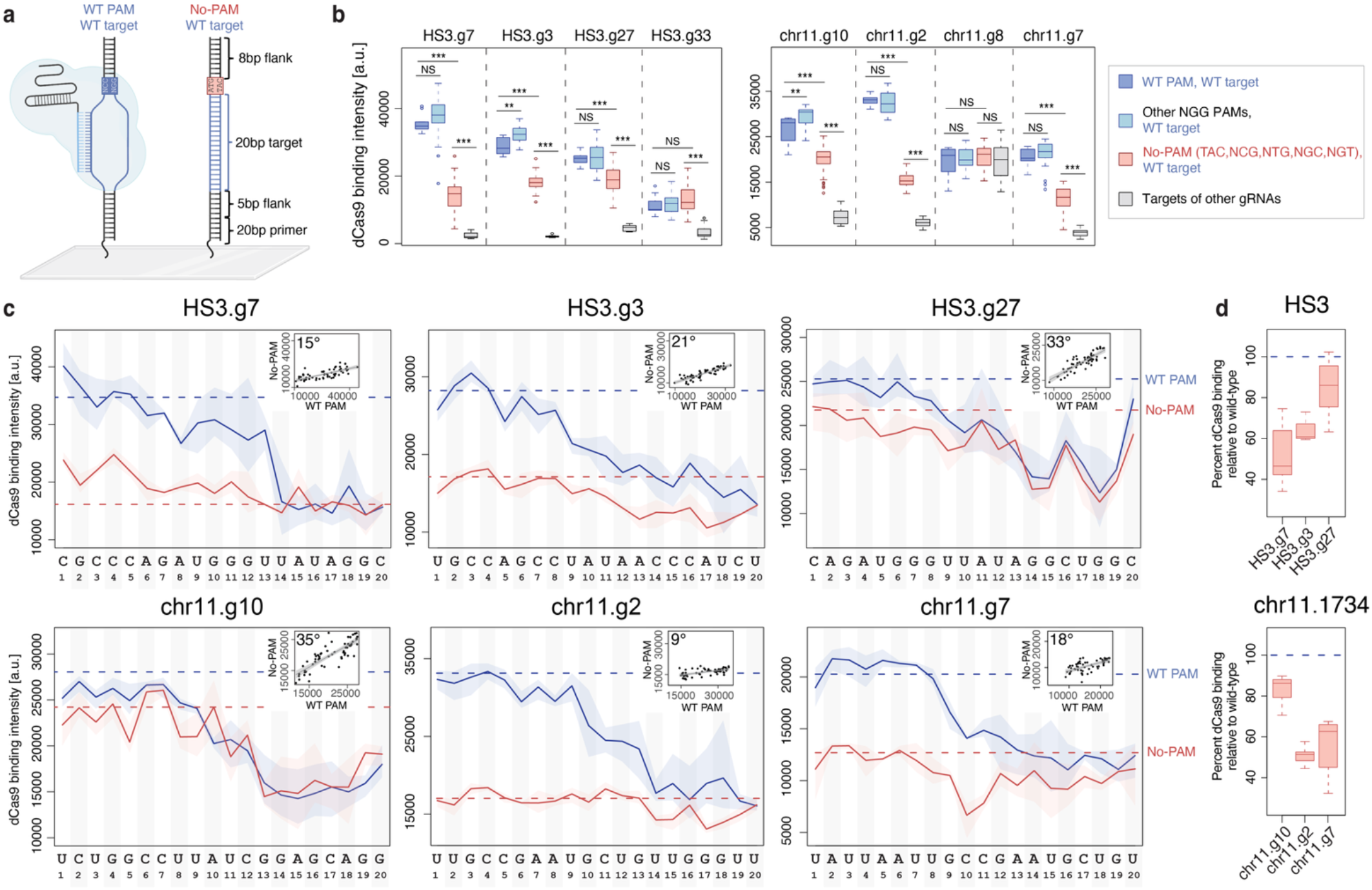
Some gRNAs show strong binding to DNA sites with extensive protospacer complementarity even in the absence of a PAM. **a**, Design of WT PAM and no-PAM probes for TANGO DNA libraries. **b**, TANGO binding intensities for HS3 and chr11.1734 CRISPRi guides on WT PAM WT-target probes (dark blue), alternative NGG PAM variants (light blue), no-PAM variants (red), and negative-control targets (i.e. targets of other, non-overalpping guides printed on the same array; gray). Statistical significance according to two-sided Mann-Whitney U-test: *P < 0.05; **P < 0.005; ***P < 0.05. **c**, TANGO mismatch tolerance profiles for HS3.g7, HS3.g3, HS3.g27, chr11.g10, chr11.g2, and chr11.g7, for both WT PAM (blue) and no-PAM (red) sequences, plotted across target site positions 1-20. Blue lines show the mean binding intensity at each position, calculated over the three possible mismatches at that position, for sequences containing the WT PAM. Red lines show the mean binding intensity at each position, calculated over the three possible mismatches at that position, for sequences where the WT PAM was replaced by TAC (i.e. no-PAM). Horizontal dotted lines show the binding level at the WT target site with either the WT PAM (blue) or the no-PAM TAC (red). Blue and red shading shows the range of binding intensities across the mismatches at each position. Insets show the correlation between the mismatch tolerance profiles for PAM vs. no-PAM probes, with the slope of the best fit line specified on the plots. A slope angle of 45° would correspond to equally strong binding to PAM and no-PAM sequences, i.e. complete PAM independence. **d**, PAM dependence summarized as the no-PAM (TAC) binding level relative to the WT PAM binding level, for the probes with WT targets sites. The distributions shown are over the replicate DNA spots for the no-PAM sequences.

When the PAM was replaced with a no-PAM sequence, the two non-working guides HS3.g33 and chr11.g8, which have little or no guide-specific DNA binding, showed minimal changes in TANGO signal (**Figure 6b**), consistent with their lack of efficient guide-directed target recognition. In contrast, all gRNAs with guide-specific DNA binding showed the expected reduction in binding to the no-PAM probes (**Figure 6b**, dark blue versus red), consistent with the established requirement of Cas9/dCas9 for a PAM for efficient target recognition^21,22,48,67^. Surprisingly, binding to no-PAM probes remained substantially higher compared to negative control probes (**Figure 6b**; red vs grey boxes, two-sided Mann-Whitney U test *p*<0.0005), indicating that elimination of the PAM at an otherwise perfect target site does not reduce binding to background levels. This behavior was observed consistently across all gRNAs with guide-specific DNA binding, including the four ChIP-benchmarking guides and the eleven guides targeting the *FOXO4* promoter (**Figure S6g**), whose TANGO libraries also contained probes in which the perfect protospacer was followed by the no-PAM trinucleotide TAC. Binding to the TAC no-PAM probes was assessed by EMSA for three of the gRNAs tested by TANGO, and they confirmed weaker but guide-specific binding to the TAC probes (**Figure S6d-f**), especially under binding conditions that included molecular crowders meant to mimic the crowded cellular environment^68–70^. Together, these results indicate that although an NGG PAM is required for maximal dCas9 binding, extensive guide-target complementarity can still promote substantial DNA binding in its absence.

Next, we asked whether the residual binding observed in the absence of a PAM reflects guide-directed DNA recognition or nonspecific interactions. To address this question, we compared mismatch tolerance profiles obtained for wild-type PAM vs. no-PAM target sequences. We focused on a representative no-PAM sequence, TAC, because neither TAC nor its reverse complement corresponds to the canonical NGG PAM or the commonly reported non-canonical NAG and NGA PAMs. The non-specific guides chr11.g8 and HS3.g33 were excluded from this analysis, as they did not exhibit PAM-dependent binding (**Figure 6b**).

Across gRNAs, mismatch tolerance profiles obtained with the wild-type PAM and the TAC no-PAM sequence were generally well correlated (**Figure 6c**; Pearson’s R² = 0.53, 0.75, and 0.72 for HS3.g7, HS3.g3, and HS3.g27, respectively, and 0.70, 0.35, and 0.41 for chr11.g10, chr11.g2, and chr11.g7), indicating that a substantial fraction of the residual no-PAM binding remains guide-directed and sequence-specific. However, the extent to which DNA binding was reduced in the absence of a PAM varied widely among guides, resulting in substantially different slopes for the relationship between the wild-type PAM and no-PAM mismatch tolerance profiles (best-fit line angles ranging from 9° to 35°, where 45° corresponds to equal binding under the two conditions; **Figure 6c**). Most gRNAs exhibited relatively flat slopes (less than 21°), indicating that high-affinity binding by those gRNAs is highly dependent on the presence of a PAM sequence. In contrast, HS3.g27 and chr11.g10 displayed 33-35° slopes, indicating that they retained a much larger fraction of their binding in the absence of a PAM (**Figure 6c**). These findings reveal substantial gRNA-to-gRNA variation in PAM dependence, a trend confirmed by mismatch tolerance analyses for the ChIP-benchmarking and the FOXO4 guides in our study (**Figure S6i**), and raise the possibility that some gRNAs may engage genomic sites with high protospacer complementarity even in the absence of a PAM. The molecular rules driving strong no-PAM recognition for some gRNAs, and to what extent such interactions occur in cells and contribute to off-target binding, remain to be determined in future studies.

### TANGO reveals effects of the downstream PAM-flanking sequence on dCas9 binding

Previous studies have suggested that the DNA sequence immediately 3′ of the PAM, outside of the protospacer, can influence Cas9 target recognition and activity, although these effects remain incompletely understood and have not been systematically characterized for dCas9 binding^15,71,72^. Here, we refer to the 8-bp region immediately 3′ of the PAM and outside the protospacer as the downstream PAM-flanking sequence. To isolate its contribution to target recognition, we used TANGO to generate, for the eight gRNAs targeting the HS3 and chr11.1734 regions, DNA probe variants in which the protospacer and PAM were held constant while systematically modifying only the downstream PAM-flanking sequence (**Figure S7a**).

First, to determine whether the nucleotide composition of the downstream PAM-flanking sequence influences binding, we generated variants containing 0%, 25%, 50%, or 75% GC (0, 2, 4, or 6 G/C bases, respectively), while maintaining the wild-type protospacer and PAM. For the 25% and 50% GC designs, we further positioned the G/C bases either adjacent to the PAM or at the distal end of the flanking region, allowing us to distinguish the effects of GC content from those of GC position (**Figure S7a**). Increasing GC content progressively reduced dCas9 binding for all seven gRNAs with guide-dependent recognition (**Figure S7b**), i.e. all gRNAs except chr11.g8, which we previously determined not to assemble efficiently with dCas9 (**Figure 5**). Furthermore, for both the 25% and 50% GC designs (i.e. 2 G/C and 4 G/C), positioning the G/C bases farther from the PAM resulted in stronger binding than placing them immediately adjacent to the PAM (**Figure S7c**). These observations suggest that elevated GC content immediately 3′ of the PAM may impede the early steps in target recognition, potentially by reducing the efficiency of local DNA strand separation required for R-loop initiation.

Next, we asked whether occurrences of PAM-like sequences downstream of the actual PAM influence binding, independently of overall GC content. To this end, we generated probes containing six G/C bases in the 3’ flanking region of the PAM, arranged to create additional PAM-like motifs (**Figure S6d,e**). Specifically, we introduced: (i) a single CCN proximal to the PAM, (ii) a single CCN distal to the PAM, (iii) both proximal and distal CCN motifs, and (iv) similar configurations of NGG motifs. Here, CCN corresponds to an NGG motif on the opposite DNA strand relative to the native PAM orientation, whereas NGG is in the same orientation as the native PAM. We found that introducing a proximal CCN motif downstream of the NGG PAM led to a slight decrease in the DNA-binding level across all gRNAs tested (**Figure S6f**). One possible explanation for this trend is that the CCN motif acts as an NGG PAM on the opposite DNA strand, thus creating a decoy PAM that transiently recruits the dCas9 RNP in a nonproductive orientation, slowing down recognition of the true on-target site. Although this reverse-orientation sampling has not, to our knowledge, been directly demonstrated, it is consistent with the current model of Cas9 target search, in which Cas9 transiently samples candidate PAM sites and initiates DNA interrogation and RNA:DNA heteroduplex formation only after PAM recognition^20,48,67^.

Surprisingly, introducing a distal CCN motif produced a substantial increase in binding for all gRNAs, with the increase being most pronounced at negative control probes (**Figure S6f**, grey boxes). This suggests that the observed effect is not due to guide-specific binding. Instead, it is likely an artifact due to the array conditions, potentially due to the positioning of the distal CCN four base pairs away from the DNA end, at a location close to where an active Cas9 RNP would cleave DNA. While we do not expect this type of binding to occur often in cells, our observation does raise questions about how dCas9 might interact with free DNA ends containing PAM-like motifs.

In contrast to CCN motifs, introducing one or two additional NGG motifs in the same orientation as the native NGG PAM had only minimal effects on DNA binding, regardless of their position within the downstream PAM-flanking sequence, and across all gRNAs tested (**Figure 7g**). Previous work showed that a high density of NGG PAMs can increase Cas9 residence time on DNA in the absence of gRNA complementarity^48^, a trend that we also observed for dCas9 in our TANGO measurements (**Figure S7h**). However, our data also indicate that once a canonical PAM and a fully complementary protospacer are present, the addition of one or two nearby NGG motifs provides little or no further enhancement of dCas9 binding.

Collectively, the results presented in this study demonstrate that TANGO enables comprehensive, quantitative dissection of the intrinsic determinants of RNP-DNA recognition. TANGO not only recapitulates established features of CRISPR target recognition, including PAM dependence and position-dependent mismatch tolerance, but also reveals additional determinants of binding, including gRNA-specific differences in mismatch tolerance and RNP assembly, and sequence features within the downstream PAM-flanking region. Compared with existing high-throughput binding assays, TANGO’s high sensitivity enables robust measurements across a broader range of binding affinities, allowing subtle yet biologically meaningful differences in target recognition to be resolved. Importantly, TANGO identifies gRNAs with intrinsically weak DNA binding, poor specificity, or defective dCas9 RNP assembly, providing a mechanistic explanation for why some gRNAs fail to elicit robust transcriptional regulation in CRISPRi experiments despite appearing suitable based on existing tools. Together, these findings establish TANGO as a versatile platform for systematically interrogating the intrinsic molecular rules governing dCas9 target recognition and binding specificity, providing a mechanistic foundation for understanding gRNA performance and for improving the design of CRISPR-based technologies for epigenome editing.

## DISCUSSION

A central challenge in CRISPRi/a gRNA design is that poor performance in cells can arise from multiple, mechanistically distinct causes: (i) inefficient dCas9 RNP assembly, (ii) weak on-target binding, (iii) excessive mismatch tolerance that dilutes activity across competing, off-target sites, or (iv) restricted cellular context limiting access to an otherwise competent target (e.g. due to inaccessible chromatin or competitive DNA binding by transcription factor proteins). Even for well characterized CRISPRi/a gRNA promoter libraries^6,33,73^, unexpected off-target binding can be misleading and confound interpretation^58^. These challenges are likely to be even greater in screens targeting distal regulatory elements, where the determinants of guide performance remain less well understood. Because multiple independent mechanisms can produce the same apparent phenotype of poor gRNA activity, there is a need for quantitative, high-throughput approaches that systematically isolate and dissect these mechanisms and distinguish intrinsic properties of the dCas9:gRNA complex from the influence of the cellular environment.

TANGO directly addresses this challenge by measuring the intrinsic interaction between dCas9:gRNA complexes and tens of thousands of defined DNA targets in parallel in a cell-free system. This provides a mechanistic framework for distinguishing gRNAs that fail because of weak target recognition, low specificity, or poor RNP assembly, from those whose activity is primarily constrained by the cellular environment. Such mechanistic resolution is essential for interpreting CRISPRi/a experiments, understanding unexpected gRNAs behavior, and should facilitate the development of improved gRNAs design algorithms and the rational engineering of increasingly precise CRISPR-based technologies. While other cell-free assays are available for characterizing gRNAs (e.g. ^28–31,47,48^), TANGO has the key advantage of combining high sensitivity with the capacity to interrogate tens of thousands of target sequences in parallel.

Across benchmark gRNAs with previously published biochemical and ChIP-seq datasets^22,28^, TANGO faithfully recapitulated canonical features of dCas9 target recognition, including gRNAs-specific binding to the native targets, strong sensitivity to PAM-proximal mismatches, and expected mismatch-identity effects. Importantly, TANGO revealed these specificity patterns even for gRNAs with relatively weak intrinsic binding, for which previous high-throughput binding studies^28^ showed little apparent positional discrimination or fell near the assay’s detection limit (**Figure 2a,b**). These results demonstrate that TANGO substantially expands the dynamic range and sensitivity with which intrinsic dCas9-DNA interactions can be quantified, enabling mechanistic characterization of guides that were previously difficult to study biochemically.

Intrinsic binding measured by TANGO also correlated with cellular occupancy, with the strongest agreement observed at targets located in accessible chromatin. This is consistent with previous studies showing that nucleosomes and higher-order chromatin structure limit Cas9 and dCas9 access to otherwise compatible DNA sequences^74,75^. As expected given the complexity of the nuclear environment, we identified numerous genomic sites that bound dCas9 strongly i*n vitro* but exhibited relatively weak ChIP-seq signal, indicating that high intrinsic binding affinity alone is insufficient for robust occupancy in cells. In contrast, we did not observe genomic sites with strong ChIP-seq signal that had weak intrinsic binding in TANGO. Collectively, these observations suggest that intrinsic affinity measured by TANGO defines an upper bound on dCas9 occupancy determined solely by DNA sequence, while chromatin accessibility and other nuclear factors determine the extent to which that binding potential is realized in living cells. By separating intrinsic DNA-binding affinity from cellular influences, TANGO can therefore distinguish sequence-intrinsic defects in target recognition from limitations imposed by chromatin or other features of the nuclear environment.

One unresolved question is how intrinsically strong targets embedded within nucleosomal DNA become occupied by dCas9 in living cells. In rapidly proliferating HEK293T cells^22^, one possibility is that dCas9 gains access transiently during DNA replication, when nucleosomes are disrupted, and subsequently prevents their reassembly, primarily at the target sites bound strongly by dCas9. Interestingly, we observed the same overall relationship, i.e. higher intrinsic binding affinity for bound sites in closed vs. open chromatin (**Figure 4g**), for g31 in primary astrocytes, which divide infrequently, suggesting that chromatin disruption during DNA replication cannot fully explain dCas9 binding in nucleosome DNA. Instead, transient nucleosome unwrapping (“breathing”), chromatin remodeling, or other dynamic fluctuations in chromatin organization^75^ may provide sufficient opportunities for dCas9 binding, although the contributions of such mechanisms remain unknown. Nevertheless, our observation that targets in closed chromatin required substantially higher intrinsic TANGO binding signals than targets in open chromatin in order to achieve detectable occupancy by ChIP-seq suggests that chromatin does not fundamentally alter the intrinsic sequence preferences of dCas9. Rather, chromatin acts as an additional energetic barrier, raising the binding-affinity threshold required for productive target engagement in cells.

TANGO also identified guides with unusually promiscuous intrinsic DNA-binding behavior. In both the benchmark ChIP-seq dataset and the *FOXO4* CRISPRa locus, the guides that displayed elevated mismatch tolerance in TANGO also exhibited broad off-target occupancy in cells (g4 and g31, respectively), indicating that their extensive genomic binding is an intrinsic property of the dCas9:gRNA complex rather than solely a consequence of chromatin context and/or genome composition. Beyond identifying promiscuous gRNAs, TANGO distinguished working from non-working CRISPRi guides and, importantly, pinpointed underlying causes of gRNA failure. Across the loci examined here, poor gRNA performance could be attributed to distinct mechanisms—including weak on-target binding, inefficient dCas9:gRNA assembly, or excessive mismatch tolerance—demonstrating that TANGO can explain why a gRNA fails, rather than simply predicting that it will. By separating the molecular determinants of gRNA activity and enabling a mechanistic diagnosis of gRNA failure, TANGO can improve interpretation of CRISPRi/a experiments. Furthermore, applying TANGO to larger collections of gRNAs will allow the identification of general principles governing RNP-DNA binding activity, and ultimately enable mechanism-informed models for CRISPRi/a gRNA optimization.

A particularly intriguing finding of this study is that TANGO revealed an additional, orthogonal dimension of gRNA specificity related to PAM dependence. Although high-affinity DNA binding was strongly dependent on a canonical PAM for most of the 23 gRNAs examined, two gRNAs (HS3.g27 and chr11.g10) retained substantial guide-specific binding even in the absence of a PAM, including when the NGG PAM was replaced by TAC, a trinucleotide expected to be highly nonpermissive for SpCas9/dCas9 binding and activity. Wild-type SpCas9 can cleave fully complementary targets with certain noncanonical PAMs, particularly NAG and NGA, although substantially less efficiently than targets with canonical NGG PAMs^19,66^. Cell-free studies have also detected cleavage at non-NGG targets, especially at high RNP concentrations^76^, but the resulting PAM motif was mainly supporting the non-canonical NAG and NGA PAMs. We are not aware of any evidence supporting efficient binding or cleavage at strongly nonpermissive sequences such as TAC. TANGO nevertheless detected substantial, above-background dCas9 binding to perfectly matched protospacers adjacent to TAC for all gRNAs tested (**Figure 6b, S6g**), with two gRNA, HS3.g27 and chr11.g10, showing a binding reduction of less than 15% when the wild-type PAM was replaced by TAC (**Figure 6d**). These two gRNAs also exhibited guide-specific mismatch tolerance profiles under no-PAM conditions, indicating that the no-PAM binding pattern does not simply reflect nonspecific interactions.

Whether such PAM-independent binding contributes to genomic off-target occupancy remains to be determined. Although PAM-independent interactions are weaker than canonical PAM-dependent interactions (**Figure 6,S6**) and are therefore unlikely to support efficient DNA cleavage by nuclease-active Cas9, they may nevertheless be functionally relevant in CRISPRi/a and other dCas9-based epigenome editing applications, where DNA binding rather than DNA cleavage underlies biological activity. Measurements in living cells indicate that dCas9 remains bound to perfectly matched targets with NGG PAMs for several hours, whereas even a single mismatch can reduce residence times to only a few minutes while still permitting stable target occupancy^77^. By comparison, most sequence-specific transcription factors occupy their binding sites for only seconds to tens of seconds, yet these transient interactions are sufficient to drive robust transcriptional regulation^78–81^. Although the residence time of dCas9 at no-PAM targets is unknown, our observation that some gRNAs retain substantial, sequence-specific DNA binding in the absence of a PAM, comparable or even higher than binding to mismatched targets with an NGG PAM (**Figure 6c,S6i**), raises the possibility that these no-PAM interactions could persist long enough to influence gene expression, despite likely being too short-lived to support efficient DNA cleavage. This possibility will require direct investigation in future studies.

The molecular basis for this gRNA-dependent PAM independence remains unknown. One possible technical explanation is that a small fraction of DNA molecules on the TANGO arrays remain single-stranded despite the primer-extension step used to generate double-stranded DNA, a strategy widely employed in high-density protein-DNA binding array platforms^36,39,41,42,82^. Given that Cas9:gRNA RNPs can bind complementary ssDNA in a PAM-independent manner^48,83,84^, such ssDNA molecules could contribute to the no-PAM binding detected in our assay. However, this explanation is unlikely to account for the large difference in no-PAM binding among gRNA, given that all gRNA were assayed on identical DNA arrays under the same experimental conditions. Moreover, our orthogonal EMSA validation further argues that the observed no-PAM binding reflects bona fide dsDNA recognition rather than binding to residual ssDNA. In these experiments, only the non-target strand was fluorescently labeled, while the target strand was unlabeled. Together, these observations argue that the no-PAM binding detected by TANGO and confirmed by EMSA cannot be explained by residual ssDNA, but instead reflects an intrinsic property of particular gRNAs rather than a general feature of the assay.

Consistent with this interpretation, the two outlier gRNA with strong PAM-independent binding share a specific sequence feature: both contain internal NGG motifs within the seed, i.e. the PAM-proximal region of the protospacer. This observation is reminiscent of the “bad seeds” identified in CRISPRa screens by Norman and colleagues^27^, who reported seed-driven effects associated with extensive off-target dCas9 occupancy. Importantly, the bad seeds were enriched for PAM-like NGG or NAG motifs, or their reverse complements (although such motifs were not present in every bad seed). Together, these observations raise the possibility that PAM-like sequences embedded within the seed can engage the PAM-interacting surface of dCas9 or otherwise facilitate the earliest steps of target interrogation. For HS3.g27 and chr11.g10, such internal motifs might act as imperfect PAM surrogates, partially compensating for the absence of a canonically positioned PAM and thereby promoting stable gRNA-directed binding. Although this model remains speculative and will require direct biochemical and structural testing, it offers a potential explanation for both the strong gRNA dependence of no-PAM binding by TANGO and the disproportionate off-target activity associated with PAM-rich seeds in CRISPRa screens. Testing whether this mechanism acts in living cells will likely require specifically designed reporters or genome-engineering experiments, as identical protospacer sequences adjacent to both canonical and strongly non-permissive PAMs are exceedingly rare in the genome (indeed, no such pair exists for any of the gRNAs examined in this study), making it difficult to isolate the contribution of PAM identity from differences in target sequence and chromatin context. More broadly, these findings suggest that PAM dependence may be influenced not only by the precise protospacer-adjacent trinucleotide but also by PAM-like sequence features embedded within the target itself, extending the repertoire of genomic sites capable of supporting stable dCas9 engagement beyond those predicted by conventional PAM-based off-target models.

Despite key advances of TANGO, several limitations should be considered. First, although the number of gRNAs profiled by TANGO is substantial, many more gRNAs spanning diverse sequence contexts will be required to derive broadly applicable principles for optimal gRNA design. Second, our mechanistic interpretations of PAM-independent binding and the effects of downstream PAM-flanking sequence context will require orthogonal biochemical and structural validation, as well validation in cellular systems. Finally, we anticipate that the TANGO framework can and should be extended to other Cas proteins and epigenome-editing architectures, to learn about the similarities and differences in intrinsic Cas-DNA recognition across systems.

In conclusion, TANGO provides a versatile framework that addresses several unmet needs across both translational and basic genome engineering communities. First, for therapeutic developers, TANGO will enable a more comprehensive evaluation of binding specificity and affinity prior to gRNA selection for clinical testing, thus reducing downstream risk in therapeutic development pipelines. Second, TANGO offers mechanistic insights into why particular gRNAs produce distinct phenotypic outcomes, allowing direct linkage between binding behavior and functional effects. This is especially relevant in light of our recent findings on widespread off-target activity of a gRNA targeting the *FOXO4* promoter^58^, which underscore the limitations of current design heuristics and highlight opportunities for improving large-scale gRNA library construction. Accordingly, TANGO should prove valuable for groups developing next-generation libraries with reduced off-target and suboptimal activity, as well as for companies engaged in gRNA synthesis and design, where empirical binding data can inform more accurate recommendations. Third, TANGO can be readily applied to profile chemically-modified gRNAs, and can be extended beyond dCas9 to other programmable nucleases including alternative CRISPR systems (e.g. dCas12a), as well as zinc-finger nucleases, TALENs, and meganucleases. Finally, beyond DNA binding, TANGO can also be expanded to independently interrogate downstream mechanistic steps in CRISPR base editing and prime editing, as well as cleavage-associated dynamics in systems employing catalytically active Cas9. Together, these applications position TANGO as a broadly enabling tool for improving both the precision and interpretability of programmable DNA targeting technologies.

## METHODS

### Selection of gRNAs for high-throughput dCas9 binding measurements

High-density DNA libraries were designed to measure dCas9 binding for 23 gRNAs spanning four experimental gRNA sets. These included: 1) four gRNAs (g1-g4) used for benchmarking against published dCas9 ChIP-seq data^22^ and high-throughput filter-binding measurements^28^; 2) eleven gRNAs targeting the *FOXO4* gene promoter and 5′ untranslated region, that were previously tested and showed activity in CRISPRa experiments in primary astrocytes^58^; 3) four gRNAs targeting the HS3 regulatory element upstream of the *HBE1* gene, with only two of four gRNAs showing activity by CRISPRi^59^; and 4) four gRNAs targeting the chr11.1734 cis-regulatory element linked to the *LMO2* gene, with only two of four gRNAs showing activity by CRISPRi^63^. The full gRNA sequences and their target sites are listed in **Table S1**. The four benchmarking gRNAs (g1-g4) were selected because they span a range of off-target binding behaviors and have both published biochemical and ChIP-seq binding data. The FOXO4 gRNA set was selected to test whether TANGO can capture DNA-binding specificity for a variety of gRNAs that are active in CRISPRa. The HS3 and chr11.1734 gRNA sets were selected because they each contained closely spaced targets for gRNAs with similar sequence-based specificity predictions (by GuideScan2^32^) but divergent CRISPRi activity, enabling direct comparison of working versus non-working gRNA within the same local genomic context.

### DNA library designs

Each arrayed DNA probe consisted of a 60-bp sequence containing a constant primer-complement region and a variable target region composed of a 20-nt protospacer sequence, a 3-nt PAM, and native genomic flanks. Each unique probe sequence was represented in 8 or 9 replicate spots (depending on the array design; **Table S1**), which we showed in previous work to be sufficient for obtaining robust and highly-reproducible data on high-density Agilent arrays^38,39,45^. For the core TANGO assay, we designed three main classes of probes for each gRNA, to be synthesized together on the DNA arrays. First, “WT target” probes contained the perfectly matched 20-nt protospacer sequence and native NGG PAM embedded in native genomic sequence context, with 5-nt upstream of the protospacer and 8-nt downstream of the PAM, with the PAM-proximal flanking region located at the free DNA end of the probe. This probe configuration was chosen because it resulted in a wide range of dCas9 binding intensities in our preliminary tests. Second, “mutated target” probes were designed to contain all possible single-nucleotide substitutions across the 20-nt protospacer (3 × 20 probes per gRNA), while keeping the PAM and flanking sequences constant. Third, no-PAM probes in which the WT PAM was replaced with 5’-TAC-3’, together with the corresponding WT target and flanks, as well as all protospacer 1-nt mismatch variants, were designed as controls for all gRNAs tested.

For the ChIP-benchmarking guides, the array also included two types of negative control probes: 1) three synthetic random sequences with varied NGG/CCN content and position, and 2) 60 random genomic fragments from open chromatin regions in HEK293T cells, lacking designed complementarity to the tested guides. The TANGO DNA library design also included off-target sequences identified from published ChIP-seq data, each embedded in its native genomic context.

For the FOXO4 gRNA set, the array included the same synthetic random sequences used as negative-control probes for the ChIP-benchmarked gRNAs.

For the CRISPRi gRNA sets (i.e. the HS3 gRNAs and the chr11.1734 gRNAs), a PAM-series was additionally designed. For each WT target, probes were synthesized bearing the native WT NGG PAM, the three alternative NGG PAM variants, and a no-PAM series in which the first base of the PAM was kept identical to the WT PAM “N” but the downstream GG was replaced with alternative dinucleotides excluding NGG, NAG, and NGA.

To examine PAM-adjacent sequence-context effects, for the CRISPRi gRNA sets, additional probe variants were designed in which the protospacer and WT NGG PAM were held constant while the 8-bp PAM-adjacent flank was systematically modified. Starting from a 0GC backbone 5’-TATAATTA-3’ with 0% GC-content, variants containing 2, 4, or 6 G/C bases were generated. For the 2GC and 4GC designs, G/C bases were positioned either PAM-proximal (p) or PAM-distal (d) within the 8-bp flank. The precise flanking sequences used were TATAATTA (0GC), T<u>GC</u>AATTA (2GCp), TATAA<u>GC</u>A (2GCd), T<u>GCGC</u>TTA (4GCp), TAT<u>GCGC</u>A (4GCd), T<u>GCGCGC</u>A (6GC), with the G/C-containing segment underlined. Additional PAM-like designs were built on the 6GC backbone by introducing CCN or NGG motifs in PAM-proximal, PAM-distal, or both positions, while keeping the overall GC content constant. The resulting flanking sequences were T<u>CGG</u>CGCA (1NGGp), TGCG<u>CGG</u>A (1NGGd), T<u>CGGCGG</u>A (2NGG), T<u>CCG</u>CGCA (1CCNp), TGCG<u>CCG</u>A (1CCNd), T<u>CCGCCG</u>A (2CCN), with the PAM-like motifs underlined. These designs were used to quantify the effect of local GC content, GC placement, and nearby PAM-like motifs on the TANGO binding readout.

### DNA array preparation

TANGO DNA libraries were synthesized on microarray slides (Agilent) in various formats: 8x60k, 16x25k, or 24x13k, which correspond to 8, 16, or 24 chambers, each containing approximately 60,000, 25,000, or 13,000 DNA spots, respectively. For each unique DNA sequence, 8 or 9 replicate spots (depending on the array format and the size of the DNA library) were randomly distributed across the microarray surface within each chamber. Each DNA sequence in the library was synthesized as a 60-nt single-stranded DNA (ssDNA) comprising a 36-nt custom, variable sequence followed by a 24-nt constant sequence (5′-GTCTGTGTTCCGTTGTCCGTGCTG-3′) that is complementary to a DNA primer used for primer extension to double-strand the DNA on the array, as described previously^42,82^. Briefly, the array was incubated in a reaction mixture containing 26 mM Tris-HCl, pH 9.5, 6.5 mM MgCl2, 1.16 uM primer, 163 uM dNTPs, 32 unit Thermo Sequenase polymerase. The incubation was then performed sequentially at 85 °C for 10 mins, 75 °C for 10 mins, 65 °C for 10 mins, and 60 °C for 90 mins.

### High-throughput on-chip RNP-DNA binding assay

The ribonucleoprotein (RNP) complex was first assembled by incubating 2uM dSpCas9-His (IDT Alt-R™ S.p. dCas9 Protein V3) with 4.8uM sgRNA (IDT Alt-R™ CRISPR Custom Guide RNAs) in the reaction buffer (20 mM Tris-HCl, pH 7.4, 150 mM KCl, 10% glycerol, 5 mM MgCl2 and 1 mM TCEP) for 10 mins at 37°C. The RNP-DNA binding assay was adapted from the universal protein-DNA binding microarray (PBM) protocol^82^. The double-stranded DNA array was first blocked with 2% nonfat milk (Sigma) for 30 mins, followed by incubation with RNP complexes for 1 hour at room temperature. RNP binding experiments were performed with either 100 nM, 50 nM, or 20 nM RNP in a binding buffer containing 20 mM HEPES-KOH, pH7.5, 150 mM KCl, 5 mM MgCl₂, 5% glycerol, 0.025% TX-100, and 1 mM TCEP. Following the incubation with RNP complexes, the array was incubated with 10 ug/mL Penta-His AlexaFluor488-conjugated antibody (Qiagen) to label the bound RNP. Binding intensity was quantified using a GenePix 4400A microarray scanner with GenePix Pro 7.0 software, as previously described^82^. For each unique sequence, the median fluorescence intensity was computed over the 8 or 9 replicate DNA spots printed with that particular sequence.

### Electrophoretic mobility shift assays

Select sequences from the TANGO DNA libraries were also tested by electrophoretic mobility shift assays (EMSAs), using protocols adapted from ^85–87^, as described below. Template DNA sequences used in EMSAs matched the design of target sequences on the TANGO arrays, including the 36-nt variable sequence and 24-nt 3’ constant sequence. A primer complementary to the constant region was designed with a 5’ IR700 modification (5’- /5IRD700/CAGCACGGACAACGGAACACAGAC-3’). All oligos were ordered as single-stranded DNA from Integrated DNA Technologies (Coralville, IA).

An extension reaction was performed on template ssDNA for each DNA sequence of interest. Triplicate master mixes of 50 μL each were prepared and split into separate wells of 8-well PCR strip tubes: 10 μL 5x Q5 buffer, 0.5 μL Q5 polymerase (New England Biolabs), 1 μL dNTPs (10 mM each) (New England Biolabs, NEB), 2.5 μL Constant-IR700 primer (10 μM), 10 μL template ssDNA (1 μM), 34 μL nuclease-free water. Master mixes were incubated in a BioRad T100 thermocycler with a primer extension program (105°C heated lid): 98°C 2 min, 72°C 3 min, 4°C forever. Following extension, dsDNA triplicate reactions were consolidated and cleaned using Monarch® Spin PCR & DNA Cleanup Kit (NEB) and its “DNA Cleanup and Concentration” protocol. Cleaned products were incubated with 15 μL of 37°C pre-warmed nuclease-free water, eluted, and quantified with Qubit™ 1X dsDNA High Sensitivity (Invitrogen).

dCas9 RNPs were assembled with 250 nM dCas9 and 375 nM sgRNA in a final volume of 10 μL. An assembly mix was prepared in 5 μL total: 1 μL 5x reaction buffer (100 mM HEPES-KOH pH 7.5, 750 mM KCl, 25 mM MgCl_2_, 50% glycerol), 0.40 μL dCas9 (6.2 μM) (IDT), 1 μL TCEP (5 mM made fresh, ThermoFisher), 0.38 μL sgRNA (10 μM) (IDT), 2.22 μL nuclease-free water. Negative controls were prepared with no dCas9 or sgRNA, substituting with nuclease-free water. The assembly mix was mixed gently by flicking and spun down, then incubated in a thermocycler (heated lid: off): 37°C 10 min, followed by 4°C.

When the assembly reaction was complete, a binding mix was prepared to yield 5 nM IR700-labeled dsDNA in a final volume of 10 μL: 1 μL 5x reaction buffer, 1 μL IR700-labeled extended dsDNA (50 nM), 0.5 μL BSA (4 mg/ml) (New England Biolabs), 0.25 μL Triton X-100 (1%) (ThermoFisher), 1 μL TCEP (5 mM), 5 μL Assembly Mix, 1.25 μL nuclease-free water. The binding mix was mixed gently by flicking and spun down, then incubated in a thermocycler (heated lid: off): 20°C 1 hour, followed by 4°C.

Next, 10 μL of binding mix was combined with 2 μL of 6x purple loading dye without SDS (NEB). A 15-well 8% Novex™ TBE PAGE gel (ThermoFisher) was rinsed, then loaded with 800 mL of 0.5x TBE running buffer (BioRad). Loading wells were flushed thoroughly with TBE running buffer. Next, 12 μL of binding mixture was loaded into each well. All PAGE gels were run in a 4°C cold room at 200V for 40 minutes, then visualized on a Li-cor Odyssey CLx imager. Bands were analyzed and quantified using ImageJ.

### ChIP-seq data processing

For the g4 ChIP-seq data from Kuscu et al.^22^, raw sequencing reads (FASTQ files) were aligned to the hg19 reference genome using Bowtie2 (*-p 8 -t -q*). Reads that were unaligned, had low mapping quality, or represented non-primary alignments were removed using SAMtools (samtools view -b -F 772 -q 24). PCR duplicates, annotated using Picard MarkDuplicates, were filtered out using SAMtools, and the resulting BAM files were used for downstream peak calling. ChIP-seq peaks were identified with MACS3^88,89^ using default parameters and a p-value cutoff of 10^-5^. As in the original study^22^, ChIP-seq data generated using Cas9 alone was used as the control for peak calling.

ChIP-seq data for the g31 RNP was generated in our recent study^58^. Briefly, primary human astrocytes (ScienCell, catalog #1800) were transduced with an all-in-one CRISPRa lentiviral plasmid expressing ^VP64^dSpCas9^VP64^ together with the gRNA scaffold for either g31 or a non-targeting gRNA. Cells were harvested 8 days after transduction, and ChIP experiments were performed in biological triplicate using monoclonal Anti-FLAG M2 antibody (Sigma F1804). Input DNA controls were generated for all replicates. ChIP-seq libraries were prepared using the KAPA HyperPrep Kit (Roche) and sequenced on an Illumina NextSeq instrument using a P4 XLEAP-SBS reagent kit with a 1% PhiX spike-in, yielding approximately 30-100 million reads per sample. Additional experimental details are provided in ^58^.

For the analyses performed in this study, g31 ChIP-seq reads were aligned to the hg38 reference genome and PCR duplicates were removed as described previously^58^. ChIP-seq peaks were called independently for each of the three biological replicates (using their corresponding input controls) and for the merged datasets (all three replicates combined) for both g31 and the non-targeting guide. Peak calling was performed with MACS3^88,89^ using the *-f BAMPE* option to use the observed paired-end fragment sizes, the *--call-summits* option to retain individual summit positions even when several small peaks were merged into wider peaks, a loose q-value threshold of 0.05 for the non-targeting guide, and a stringent q-value cutoff of 10^-5^ for g31.

To identify highly reproducible g31 binding events, summit-centered peaks from the merged g31 dataset were expanded by 150 bp in each direction (301 bp total) and intersected with peak summits identified in each individual g31 replicate. Only merged peaks whose expanded regions overlapped a summit in all three replicates were retained, yielding 56,250 reproducible peaks. We next removed peaks overlapping hg38 ENCODE Blacklist V2 regions^90^, as well as peaks overlapping non-targeting (NT) guide ChIP-seq peaks identified from the merged control dataset, when the g31-to-NT read pileup ratio was less than five. This filtering procedure yielded a final set of 44,732 high-confidence, highly reproducible g31-specific ChIP-seq peaks (**Table S3**). We note that this set of peaks differs slightly from that analyzed in our related study^58^, which used the standard MACS2^88^ and IDR^91^ pipeline (IDR threshold = 0.001). We applied a different analysis pipeline in our current study because the conventional MACS2-IDR workflow frequently merged nearby dCas9 binding events into broad peaks. Although this behavior did not affect the analyses in our previous study^58^, the present work requires individual binding events to be resolved as accurately as possible in order to relate *in vivo* dCas9 occupancy measured by ChIP-seq to intrinsic DNA-binding specificity measured by TANGO.

### FOXO4 g31 ChIP-seq data analyses

g31 ChIP-seq peaks were scanned on both DNA strands for occurrences of 8-bp seed sequences immediately followed by an NGG PAM. Peaks were first searched for the perfect g31 seed sequence (the PAM-proximal 8-mer, CAGGAGAG). Peaks lacking a perfect seed were then searched sequentially for seed sequences containing one, two, three, and finally four mismatches relative to the perfect seed. Each peak was assigned to the lowest-mismatch category detected, and the resulting distributions are shown in **Figure 4b**. Most subsequent analyses focused on peaks containing seeds with 0-2 mismatches. ChIP-seq peaks were also intersected with DNA accessibility peaks obtained by ATAC-seq in astrocyte cells transduced with ^VP64^dSpCas9^VP64^ and non-targeting guide, as identified in ^58^.

The distances between seed matches and ChIP-seq peak summits (**Figure 4c**) were calculated by first identifying all 0-2 mismatch seed sequences immediately followed by an NGG PAM, on either DNA strand. Each seed match was then extended to the corresponding 23-bp target site (20-bp protospacer plus 3-bp NGG PAM), and the distance between the edge of the 23-bp target site and the ChIP-seq peak summit was calculated. Distances were assigned a value of zero when the peak summit fell within the 23-bp target site.

Enrichment of 8-bp seed sequences was evaluated by comparing their frequencies within ChIP-seq peaks to those in flanking control regions. For each 301-bp peak, 301-bp flanking regions immediately upstream and downstream were extracted (**Figure 4e**). For analyses restricted to the ±100-bp or ±50-bp windows around the peak summit, the corresponding control regions were defined as the central 201-bp or 101-bp segments of the upstream and downstream flanks. Enrichment of individual seed sequences or groups of seeds was quantified using odds ratios and assessed for statistical significance with Fisher’s exact test (**Table S3**).

For comparisons between ChIP-seq and TANGO, we used the read pileup counts for the ChIP-seq peaks, and TANGO binding intensities (measured or estimated) for 8-mer seeds with 0-2 mismatches that occurred within the peaks. Because many ChIP-seq peaks contained multiple occurrences of 0-2 mismatch seeds followed by an NGG, making it ambiguous which seed represented the true binding site, this analysis was restricted to the 25,148 peaks containing a single such seed. For increased confidence that the seed in each peak corresponded to the true binding event, we only used peaks where the seed occurred within ±100-bp of the peak summit. For 0- and 1-mismatch seeds, experimentally measured TANGO binding intensities, for the seed in the genomic sequence context of the on-target site, were used. For 2-mismatch seeds, binding intensities were estimated by subtracting the penalties associated with each mismatch from the TANGO intensity of the on-target site, assuming additive mismatch effects.

## ACKNOWLEDGEMENTS

This work was supported by the National Institutes of Health (NIH) grants RM1-HG011123 (to RG, GEC, CAG), R01-MH125236 (to RG, GEC, CAG), and UM1-HG012053 (to GEC, CAG).

## DATA AVAILABILITY

The TANGO data supporting the findings in this study are available as Supplementary Tables, in Excel format. The raw TANGO data has been deposited in the Gene Expression Omnibus (GEO) and will be make publicly available upon publication. The FOXO4 g31 ChIP-seq data are available through NCBI GEO with accession number GSE311472.

## Notes

### Competing Interest Statement

CAG is a co-founder of Tune Therapeutics, Locus Biosciences, and Sollus Therapeutics, and an advisor to Sarepta Therapeutics and Pappas Capital. WZ, MT, SJR, CAG, GEC, and RG are inventors on patents or patent applications related to CRISPR epigenome editing and screening technologies.

## REFERENCES

1 Gilbert, L. A., Larson, M. H., Morsut, L., Liu, Z., Brar, G. A., Torres, S. E., Stern-Ginossar, N., Brandman, O., Whitehead, E. H., Doudna, J. A., Lim, W. A., Weissman, J. S. & Qi, L. S. CRISPR-mediated modular RNA-guided regulation of transcription in eukaryotes. Cell 154, 442–451 (2013). 10.1016/j.cell.2013.06.044

2 Qi, L. S., Larson, M. H., Gilbert, L. A., Doudna, J. A., Weissman, J. S., Arkin, A. P. & Lim, W. A. Repurposing CRISPR as an RNA-guided platform for sequence-specific control of gene expression. Cell 152, 1173–1183 (2013). 10.1016/j.cell.2013.02.022

3 McCutcheon, S. R., Rohm, D., Iglesias, N. & Gersbach, C. A. Epigenome editing technologies for discovery and medicine. Nature biotechnology 42, 1199–1217 (2024). 10.1038/s41587-024-02320-1

4 Konermann, S., Brigham, M. D., Trevino, A. E., Joung, J., Abudayyeh, O. O., Barcena, C., Hsu, P. D., Habib, N., Gootenberg, J. S., Nishimasu, H., Nureki, O. & Zhang, F. Genome-scale transcriptional activation by an engineered CRISPR-Cas9 complex. Nature 517, 583–588 (2015). 10.1038/nature14136

5 Chavez, A., Scheiman, J., Vora, S., Pruitt, B. W., Tuttle, M., E, P. R. I., Lin, S., Kiani, S., Guzman, C. D., Wiegand, D. J., Ter-Ovanesyan, D., Braff, J. L., Davidsohn, N., Housden, B. E., Perrimon, N., Weiss, R., Aach, J., Collins, J. J. & Church, G. M. Highly efficient Cas9-mediated transcriptional programming. Nat Methods 12, 326–328 (2015). 10.1038/nmeth.3312

6 Gilbert, L. A., Horlbeck, M. A., Adamson, B., Villalta, J. E., Chen, Y., Whitehead, E. H., Guimaraes, C., Panning, B., Ploegh, H. L., Bassik, M. C., Qi, L. S., Kampmann, M. & Weissman, J. S. Genome-Scale CRISPR-Mediated Control of Gene Repression and Activation. Cell 159, 647–661 (2014). 10.1016/j.cell.2014.09.029

7 Thakore, P. I., D’Ippolito, A. M., Song, L., Safi, A., Shivakumar, N. K., Kabadi, A. M., Reddy, T. E., Crawford, G. E. & Gersbach, C. A. Highly specific epigenome editing by CRISPR-Cas9 repressors for silencing of distal regulatory elements. Nat Methods 12, 1143–1149 (2015). 10.1038/nmeth.3630

8 Hilton, I. B., D’Ippolito, A. M., Vockley, C. M., Thakore, P. I., Crawford, G. E., Reddy, T. E. & Gersbach, C. A. Epigenome editing by a CRISPR-Cas9-based acetyltransferase activates genes from promoters and enhancers. Nature biotechnology 33, 510–517 (2015). 10.1038/nbt.3199

9 Yamagata, T., Raveau, M., Kobayashi, K., Miyamoto, H., Tatsukawa, T., Ogiwara, I., Itohara, S., Hensch, T. K. & Yamakawa, K. CRISPR/dCas9-based Scn1a gene activation in inhibitory neurons ameliorates epileptic and behavioral phenotypes of Dravet syndrome model mice. Neurobiol Dis 141, 104954 (2020). 10.1016/j.nbd.2020.104954

10 Policarpi, C., Munafo, M., Tsagkris, S., Carlini, V. & Hackett, J. A. Systematic epigenome editing captures the context-dependent instructive function of chromatin modifications. Nat Genet 56, 1168–1180 (2024). 10.1038/s41588-024-01706-w

11 Nunez, J. K., Chen, J., Pommier, G. C., Cogan, J. Z., Replogle, J. M., Adriaens, C., Ramadoss, G. N., Shi, Q., Hung, K. L., Samelson, A. J., Pogson, A. N., Kim, J. Y. S., Chung, A., Leonetti, M. D., Chang, H. Y., Kampmann, M., Bernstein, B. E., Hovestadt, V., Gilbert, L. A. & Weissman, J. S. Genome-wide programmable transcriptional memory by CRISPR-based epigenome editing. Cell 184, 2503–2519 e2517 (2021). 10.1016/j.cell.2021.03.025

12 Liu, X. S., Wu, H., Ji, X., Stelzer, Y., Wu, X., Czauderna, S., Shu, J., Dadon, D., Young, R. A. & Jaenisch, R. Editing DNA Methylation in the Mammalian Genome. Cell 167, 233–247 e217 (2016). 10.1016/j.cell.2016.08.056

13 Sapozhnikov, D. M. & Szyf, M. Unraveling the functional role of DNA demethylation at specific promoters by targeted steric blockage of DNA methyltransferase with CRISPR/dCas9. Nat Commun 12, 5711 (2021). 10.1038/s41467-021-25991-9

14 Chen, B., Gilbert, L. A., Cimini, B. A., Schnitzbauer, J., Zhang, W., Li, G. W., Park, J., Blackburn, E. H., Weissman, J. S., Qi, L. S. & Huang, B. Dynamic imaging of genomic loci in living human cells by an optimized CRISPR/Cas system. Cell 155, 1479–1491 (2013). 10.1016/j.cell.2013.12.001

15 Doench, J. G., Hartenian, E., Graham, D. B., Tothova, Z., Hegde, M., Smith, I., Sullender, M., Ebert, B. L., Xavier, R. J. & Root, D. E. Rational design of highly active sgRNAs for CRISPR-Cas9-mediated gene inactivation. Nature biotechnology 32, 1262–1267 (2014). 10.1038/nbt.3026

16 Doench, J. G., Fusi, N., Sullender, M., Hegde, M., Vaimberg, E. W., Donovan, K. F., Smith, I., Tothova, Z., Wilen, C., Orchard, R., Virgin, H. W., Listgarten, J. & Root, D. E. Optimized sgRNA design to maximize activity and minimize off-target effects of CRISPR-Cas9. Nature biotechnology 34, 184–191 (2016). 10.1038/nbt.3437

17 DeWeirdt, P. C., McGee, A. V., Zheng, F., Nwolah, I., Hegde, M. & Doench, J. G. Accounting for small variations in the tracrRNA sequence improves sgRNA activity predictions for CRISPR screening. Nat Commun 13, 5255 (2022). 10.1038/s41467-022-33024-2

18 Chuai, G., Ma, H., Yan, J., Chen, M., Hong, N., Xue, D., Zhou, C., Zhu, C., Chen, K., Duan, B., Gu, F., Qu, S., Huang, D., Wei, J. & Liu, Q. DeepCRISPR: optimized CRISPR guide RNA design by deep learning. Genome Biol 19, 80 (2018). 10.1186/s13059-018-1459-4

19 Hsu, P. D., Scott, D. A., Weinstein, J. A., Ran, F. A., Konermann, S., Agarwala, V., Li, Y., Fine, E. J., Wu, X., Shalem, O., Cradick, T. J., Marraffini, L. A., Bao, G. & Zhang, F. DNA targeting specificity of RNA-guided Cas9 nucleases. Nature biotechnology 31, 827–832 (2013). 10.1038/nbt.2647

20 Sternberg, S. H., LaFrance, B., Kaplan, M. & Doudna, J. A. Conformational control of DNA target cleavage by CRISPR-Cas9. Nature 527, 110–113 (2015). 10.1038/nature15544

21 Wu, X., Scott, D. A., Kriz, A. J., Chiu, A. C., Hsu, P. D., Dadon, D. B., Cheng, A. W., Trevino, A. E., Konermann, S., Chen, S., Jaenisch, R., Zhang, F. & Sharp, P. A. Genome-wide binding of the CRISPR endonuclease Cas9 in mammalian cells. Nature biotechnology 32, 670–676 (2014). 10.1038/nbt.2889

22 Kuscu, C., Arslan, S., Singh, R., Thorpe, J. & Adli, M. Genome-wide analysis reveals characteristics of off-target sites bound by the Cas9 endonuclease. Nature biotechnology 32, 677–683 (2014). 10.1038/nbt.2916

23 O’Geen, H., Henry, I. M., Bhakta, M. S., Meckler, J. F. & Segal, D. J. A genome-wide analysis of Cas9 binding specificity using ChIP-seq and targeted sequence capture. Nucleic acids research 43, 3389–3404 (2015). 10.1093/nar/gkv137

24 Johnson, D. S., Mortazavi, A., Myers, R. M. & Wold, B. Genome-wide mapping of in vivo protein-DNA interactions. Science 316, 1497–1502 (2007). 10.1126/science.1141319

25 Kaya-Okur, H. S., Wu, S. J., Codomo, C. A., Pledger, E. S., Bryson, T. D., Henikoff, J. G., Ahmad, K. & Henikoff, S. CUT&Tag for efficient epigenomic profiling of small samples and single cells. Nat Commun 10, 1930 (2019). 10.1038/s41467-019-09982-5

26 Skene, P. J. & Henikoff, S. An efficient targeted nuclease strategy for high-resolution mapping of DNA binding sites. Elife 6 (2017). 10.7554/eLife.21856

27 Southard, K. M., Ardy, R. C., Tang, A., O’Sullivan, D. D., Metzner, E., Guruvayurappan, K. & Norman, T. M. Comprehensive transcription factor perturbations recapitulate fibroblast transcriptional states. Nat Genet 57, 2323–2334 (2025). 10.1038/s41588-025-02284-1

28 Boyle, E. A., Becker, W. R., Bai, H. B., Chen, J. S., Doudna, J. A. & Greenleaf, W. J. Quantification of Cas9 binding and cleavage across diverse guide sequences maps landscapes of target engagement. Sci Adv 7 (2021). 10.1126/sciadv.abe5496

29 Zhang, L., Rube, H. T., Vakulskas, C. A., Behlke, M. A., Bussemaker, H. J. & Pufall, M. A. Systematic in vitro profiling of off-target affinity, cleavage and efficiency for CRISPR enzymes. Nucleic acids research 48, 5037–5053 (2020). 10.1093/nar/gkaa231

30 Boyle, E. A., Andreasson, J. O. L., Chircus, L. M., Sternberg, S. H., Wu, M. J., Guegler, C. K., Doudna, J. A. & Greenleaf, W. J. High-throughput biochemical profiling reveals sequence determinants of dCas9 off-target binding and unbinding. Proc Natl Acad Sci U S A 114, 5461–5466 (2017). 10.1073/pnas.1700557114

31 Jones, S. K., Jr., Hawkins, J. A., Johnson, N. V., Jung, C., Hu, K., Rybarski, J. R., Chen, J. S., Doudna, J. A., Press, W. H. & Finkelstein, I. J. Massively parallel kinetic profiling of natural and engineered CRISPR nucleases. Nature biotechnology 39, 84–93 (2021). 10.1038/s41587-020-0646-5

32 Schmidt, H., Zhang, M., Chakarov, D., Bansal, V., Mourelatos, H., Sanchez-Rivera, F. J., Lowe, S. W., Ventura, A., Leslie, C. S. & Pritykin, Y. Genome-wide CRISPR guide RNA design and specificity analysis with GuideScan2. Genome Biol 26, 41 (2025). 10.1186/s13059-025-03488-8

33 Sanson, K. R., Hanna, R. E., Hegde, M., Donovan, K. F., Strand, C., Sullender, M. E., Vaimberg, E. W., Goodale, A., Root, D. E., Piccioni, F. & Doench, J. G. Optimized libraries for CRISPR-Cas9 genetic screens with multiple modalities. Nat Commun 9, 5416 (2018). 10.1038/s41467-018-07901-8

34 IDT. CRISPR-Cas9 guide RNA design checker, <https://www.idtdna.com/site/order/designtool/index/CRISPR_SEQUENCE> (

35 Martin, V., Zhuang, F., Zhang, Y., Pinheiro, K. & Gordan, R. High-throughput data and modeling reveal insights into the mechanisms of cooperative DNA-binding by transcription factor proteins. Nucleic acids research 51, 11600–11612 (2023). 10.1093/nar/gkad872

36 Mielko, Z., Zhang, Y., Sahay, H., Liu, Y., Schaich, M. A., Schnable, B., Morrison, A. M., Burdinski, D., Adar, S., Pufall, M., Van Houten, B., Gordan, R. & Afek, A. UV irradiation remodels the specificity landscape of transcription factors. Proc Natl Acad Sci U S A 120, e2217422120 (2023). 10.1073/pnas.2217422120

37 Zhang, Y., Ho, T. D., Buchler, N. E. & Gordan, R. Competition for DNA binding between paralogous transcription factors determines their genomic occupancy and regulatory functions. Genome Res 31, 1216–1229 (2021). 10.1101/gr.275145.120

38 Afek, A., Shi, H., Rangadurai, A., Sahay, H., Senitzki, A., Xhani, S., Fang, M., Salinas, R., Mielko, Z., Pufall, M. A., Poon, G. M. K., Haran, T. E., Schumacher, M. A., Al-Hashimi, H. M. & Gordan, R. DNA mismatches reveal conformational penalties in protein-DNA recognition. Nature 587, 291–296 (2020). 10.1038/s41586-020-2843-2

39 Shen, N., Zhao, J., Schipper, J. L., Zhang, Y., Bepler, T., Leehr, D., Bradley, J., Horton, J., Lapp, H. & Gordan, R. Divergence in DNA Specificity among Paralogous Transcription Factors Contributes to Their Differential In Vivo Binding. Cell Syst 6, 470–483 e478 (2018). 10.1016/j.cels.2018.02.009

40 Barrera, L. A., Vedenko, A., Kurland, J. V., Rogers, J. M., Gisselbrecht, S. S., Rossin, E. J., Woodard, J., Mariani, L., Kock, K. H., Inukai, S., Siggers, T., Shokri, L., Gordan, R., Sahni, N., Cotsapas, C., Hao, T., Yi, S., Kellis, M., Daly, M. J., Vidal, M., Hill, D. E. & Bulyk, M. L. Survey of variation in human transcription factors reveals prevalent DNA binding changes. Science 351, 1450–1454 (2016). 10.1126/science.aad2257

41 Badis, G., Berger, M. F., Philippakis, A. A., Talukder, S., Gehrke, A. R., Jaeger, S. A., Chan, E. T., Metzler, G., Vedenko, A., Chen, X., Kuznetsov, H., Wang, C. F., Coburn, D., Newburger, D. E., Morris, Q., Hughes, T. R. & Bulyk, M. L. Diversity and complexity in DNA recognition by transcription factors. Science 324, 1720–1723 (2009). 10.1126/science.1162327

42 Berger, M. F., Philippakis, A. A., Qureshi, A. M., He, F. S., Estep, P. W., 3rd & Bulyk, M. L. Compact, universal DNA microarrays to comprehensively determine transcription-factor binding site specificities. Nature biotechnology 24, 1429–1435 (2006). 10.1038/nbt1246

43 Siggers, T., Duyzend, M. H., Reddy, J., Khan, S. & Bulyk, M. L. Non-DNA-binding cofactors enhance DNA-binding specificity of a transcriptional regulatory complex. Mol Syst Biol 7, 555 (2011). 10.1038/msb.2011.89

44 Siggers, T., Chang, A. B., Teixeira, A., Wong, D., Williams, K. J., Ahmed, B., Ragoussis, J., Udalova, I. A., Smale, S. T. & Bulyk, M. L. Principles of dimer-specific gene regulation revealed by a comprehensive characterization of NF-kappaB family DNA binding. Nat Immunol 13, 95–102 (2011). 10.1038/ni.2151

45 Zhu, W., Zhang, Y., Sahay, H., Wasserman, H., Afek, A., Williams, J., Shaltz, S., Johnson, C., Pinheiro, K., MacAlpine, D. M., Weninger, K. R., Erie, D. A., Jinks-Robertson, S. & Gordan, R. DNA mutagenesis driven by transcription factor competition with mismatch repair. Cell 188, 5735–5747 e5715 (2025). 10.1016/j.cell.2025.07.003

46 Consortium, E. P. An integrated encyclopedia of DNA elements in the human genome. Nature 489, 57–74 (2012). 10.1038/nature11247

47 Josephs, E. A., Kocak, D. D., Fitzgibbon, C. J., McMenemy, J., Gersbach, C. A. & Marszalek, P. E. Structure and specificity of the RNA-guided endonuclease Cas9 during DNA interrogation, target binding and cleavage. Nucleic acids research 43, 8924–8941 (2015). 10.1093/nar/gkv892

48 Sternberg, S. H., Redding, S., Jinek, M., Greene, E. C. & Doudna, J. A. DNA interrogation by the CRISPR RNA-guided endonuclease Cas9. Nature 507, 62–67 (2014). 10.1038/nature13011

49 Singh, D., Sternberg, S. H., Fei, J., Doudna, J. A. & Ha, T. Real-time observation of DNA recognition and rejection by the RNA-guided endonuclease Cas9. Nat Commun 7, 12778 (2016). 10.1038/ncomms12778

50 Jost, M., Santos, D. A., Saunders, R. A., Horlbeck, M. A., Hawkins, J. S., Scaria, S. M., Norman, T. M., Hussmann, J. A., Liem, C. R., Gross, C. A. & Weissman, J. S. Titrating gene expression using libraries of systematically attenuated CRISPR guide RNAs. Nature biotechnology 38, 355–364 (2020). 10.1038/s41587-019-0387-5

51 Watkins, N. E., Jr., Kennelly, W. J., Tsay, M. J., Tuin, A., Swenson, L., Lee, H. R., Morosyuk, S., Hicks, D. A. & Santalucia, J., Jr. Thermodynamic contributions of single internal rA.dA, rC.dC, rG.dG and rU.dT mismatches in RNA/DNA duplexes. Nucleic acids research 39, 1894–1902 (2011). 10.1093/nar/gkq905

52 Pacesa, M., Lin, C. H., Clery, A., Saha, A., Arantes, P. R., Bargsten, K., Irby, M. J., Allain, F. H., Palermo, G., Cameron, P., Donohoue, P. D. & Jinek, M. Structural basis for Cas9 off-target activity. Cell 185, 4067–4081 e4021 (2022). 10.1016/j.cell.2022.09.026

53 Sugimoto, N., Nakano, M. & Nakano, S. Thermodynamics-structure relationship of single mismatches in RNA/DNA duplexes. Biochemistry 39, 11270–11281 (2000). 10.1021/bi000819p

54 Manandhar, D., Song, L., Kabadi, A., Kwon, J. B., Edsall, L. E., Ehrlich, M., Tsumagari, K., Gersbach, C. A., Crawford, G. E. & Gordan, R. Incomplete MyoD-induced transdifferentiation is associated with chromatin remodeling deficiencies. Nucleic acids research 45, 11684–11699 (2017). 10.1093/nar/gkx773

55 Meers, M. P., Janssens, D. H. & Henikoff, S. Pioneer Factor-Nucleosome Binding Events during Differentiation Are Motif Encoded. Mol Cell 75, 562–575 e565 (2019). 10.1016/j.molcel.2019.05.025

56 Feng, H., Guo, J., Wang, T., Zhang, C. & Xing, X. H. Guide-target mismatch effects on dCas9-sgRNA binding activity in living bacterial cells. Nucleic acids research 49, 1263–1277 (2021). 10.1093/nar/gkaa1295

57 Reisman, S. J., Halabi, D., Miller, S. E., Song, L., Geraghty, S., Sangvai, N., Rice, G., Safi, A., Crawford, G. E. & Gersbach, C. A. Comprehensive profiling of transcription factors for reprogramming human astrocytes to neuronal cells through endogenous CRISPR-based gene activation. bioRxiv (2025). 10.1101/2025.10.11.681828

58 Reisman, S. J., Zhu, W., Miller, S. E., Halabi, D., Sangvai, N., Crawford, G. E., Gordan, R. & Gersbach, C. A. Mismatch tolerance of a gRNA for CRISPR-based gene activation confers broad activity critical for cell reprogramming. bioRxiv (2026). 10.64898/2026.02.01.703129

59 Klann, T. S., Black, J. B., Chellappan, M., Safi, A., Song, L., Hilton, I. B., Crawford, G. E., Reddy, T. E. & Gersbach, C. A. CRISPR-Cas9 epigenome editing enables high-throughput screening for functional regulatory elements in the human genome. Nature biotechnology 35, 561–568 (2017). 10.1038/nbt.3853

60 Perez-Pinera, P., Kocak, D. D., Vockley, C. M., Adler, A. F., Kabadi, A. M., Polstein, L. R., Thakore, P. I., Glass, K. A., Ousterout, D. G., Leong, K. W., Guilak, F., Crawford, G. E., Reddy, T. E. & Gersbach, C. A. RNA-guided gene activation by CRISPR-Cas9-based transcription factors. Nat Methods 10, 973–976 (2013). 10.1038/nmeth.2600

61 Cosgrove, B. D., Bounds, L. R., Taylor, C. K., Su, A. L., Rizzo, A. J., Barrera, A., Sun, T., Safi, A., Song, L., Whitlow, T., Tata, A., Iglesias, N., Diao, Y., Tata, P. R., Hoffman, B. D., Crawford, G. E. & Gersbach, C. A. Mechanosensitive genomic enhancers potentiate the cellular response to matrix stiffness. Science 390, eadl1988 (2025). 10.1126/science.adl1988

62 Rohm, D., Black, J. B., McCutcheon, S. R., Barrera, A., Berry, S. S., Morone, D. J., Nuttle, X., de Esch, C. E., Tai, D. J. C., Talkowski, M. E., Iglesias, N. & Gersbach, C. A. Activation of the imprinted Prader-Willi syndrome locus by CRISPR-based epigenome editing. Cell Genom 5, 100770 (2025). 10.1016/j.xgen.2025.100770

63 ter Weele, M., Klann, T. S., Barrera, A., Ettyreddy, A. R., Liu, S., Rickels, R. A., Bryois, J., Jiang, S., Adkar, S. S., Iglesias, N., Sullivan, P. F., Reddy, T. E., Allen, A. S., Crawford, G. E. & Gersbach, C. A. Genome-wide annotation of gene regulatory elements linked to cell fitness. BioRxiv Preprint (2025). doi: 10.1101/2021.03.08.434470

64 Kleinstiver, B. P., Prew, M. S., Tsai, S. Q., Topkar, V. V., Nguyen, N. T., Zheng, Z., Gonzales, A. P., Li, Z., Peterson, R. T., Yeh, J. R., Aryee, M. J. & Joung, J. K. Engineered CRISPR-Cas9 nucleases with altered PAM specificities. Nature 523, 481–485 (2015). 10.1038/nature14592

65 Kim, H. K., Lee, S., Kim, Y., Park, J., Min, S., Choi, J. W., Huang, T. P., Yoon, S., Liu, D. R. & Kim, H. H. High-throughput analysis of the activities of xCas9, SpCas9-NG and SpCas9 at matched and mismatched target sequences in human cells. Nat Biomed Eng 4, 111–124 (2020). 10.1038/s41551-019-0505-1

66 Zhang, Y., Ge, X., Yang, F., Zhang, L., Zheng, J., Tan, X., Jin, Z. B., Qu, J. & Gu, F. Comparison of non-canonical PAMs for CRISPR/Cas9-mediated DNA cleavage in human cells. Sci Rep 4, 5405 (2014). 10.1038/srep05405

67 Anders, C., Niewoehner, O., Duerst, A. & Jinek, M. Structural basis of PAM-dependent target DNA recognition by the Cas9 endonuclease. Nature 513, 569–573 (2014). 10.1038/nature13579

68 Collette, D., Dunlap, D. & Finzi, L. Macromolecular Crowding and DNA: Bridging the Gap between In Vitro and In Vivo. Int J Mol Sci 24 (2023). 10.3390/ijms242417502

69 Huang, J. H. & Ferrell, J. E., Jr. How does cytoplasmic crowding affect reaction rates? Mol Cell 86, 9–23 (2026). 10.1016/j.molcel.2025.12.007

70 Zhou, H. X., Rivas, G. & Minton, A. P. Macromolecular crowding and confinement: biochemical, biophysical, and potential physiological consequences. Annu Rev Biophys 37, 375–397 (2008). 10.1146/annurev.biophys.37.032807.125817

71 Corsi, G. I., Qu, K., Alkan, F., Pan, X., Luo, Y. & Gorodkin, J. CRISPR/Cas9 gRNA activity depends on free energy changes and on the target PAM context. Nat Commun 13, 3006 (2022). 10.1038/s41467-022-30515-0

72 Zhang, Q., Wen, F., Zhang, S., Jin, J., Bi, L., Lu, Y., Li, M., Xi, X. G., Huang, X., Shen, B. & Sun, B. The post-PAM interaction of RNA-guided spCas9 with DNA dictates its target binding and dissociation. Sci Adv 5, eaaw9807 (2019). 10.1126/sciadv.aaw9807

73 Horlbeck, M. A., Gilbert, L. A., Villalta, J. E., Adamson, B., Pak, R. A., Chen, Y., Fields, A. P., Park, C. Y., Corn, J. E., Kampmann, M. & Weissman, J. S. Compact and highly active next-generation libraries for CRISPR-mediated gene repression and activation. Elife 5 (2016). 10.7554/eLife.19760

74 Horlbeck, M. A., Witkowsky, L. B., Guglielmi, B., Replogle, J. M., Gilbert, L. A., Villalta, J. E., Torigoe, S. E., Tjian, R. & Weissman, J. S. Nucleosomes impede Cas9 access to DNA in vivo and in vitro. Elife 5 (2016). 10.7554/eLife.12677

75 Isaac, R. S., Jiang, F., Doudna, J. A., Lim, W. A., Narlikar, G. J. & Almeida, R. Nucleosome breathing and remodeling constrain CRISPR-Cas9 function. Elife 5 (2016). 10.7554/eLife.13450

76 Karvelis, T., Gasiunas, G., Young, J., Bigelyte, G., Silanskas, A., Cigan, M. & Siksnys, V. Rapid characterization of CRISPR-Cas9 protospacer adjacent motif sequence elements. Genome Biol 16, 253 (2015). 10.1186/s13059-015-0818-7

77 Ma, H., Tu, L. C., Naseri, A., Huisman, M., Zhang, S., Grunwald, D. & Pederson, T. CRISPR-Cas9 nuclear dynamics and target recognition in living cells. J Cell Biol 214, 529–537 (2016). 10.1083/jcb.201604115

78 de Jonge, W. J., Patel, H. P., Meeussen, J. V. W. & Lenstra, T. L. Following the tracks: How transcription factor binding dynamics control transcription. Biophys J 121, 1583–1592 (2022). 10.1016/j.bpj.2022.03.026

79 Lu, F. & Lionnet, T. Transcription Factor Dynamics. Cold Spring Harb Perspect Biol 13 (2021). 10.1101/cshperspect.a040949

80 Hager, G. L., McNally, J. G. & Misteli, T. Transcription dynamics. Mol Cell 35, 741–753 (2009). 10.1016/j.molcel.2009.09.005

81 Lionnet, T. & Wu, C. Single-molecule tracking of transcription protein dynamics in living cells: seeing is believing, but what are we seeing? Curr Opin Genet Dev 67, 94–102 (2021). 10.1016/j.gde.2020.12.001

82 Berger, M. F. & Bulyk, M. L. Universal protein-binding microarrays for the comprehensive characterization of the DNA-binding specificities of transcription factors. Nat Protoc 4, 393–411 (2009). 10.1038/nprot.2008.195

83 Gasiunas, G., Barrangou, R., Horvath, P. & Siksnys, V. Cas9-crRNA ribonucleoprotein complex mediates specific DNA cleavage for adaptive immunity in bacteria. Proc Natl Acad Sci U S A 109, E2579–2586 (2012). 10.1073/pnas.1208507109

84 Ma, E., Harrington, L. B., O’Connell, M. R., Zhou, K. & Doudna, J. A. Single-Stranded DNA Cleavage by Divergent CRISPR-Cas9 Enzymes. Mol Cell 60, 398–407 (2015). 10.1016/j.molcel.2015.10.030

85 Ryder, S. P., Recht, M. I. & Williamson, J. R. Quantitative analysis of protein-RNA interactions by gel mobility shift. Methods Mol Biol 488, 99–115 (2008). 10.1007/978-1-60327-475-3_7

86 Anders, C., Niewoehner, O. & Jinek, M. In Vitro Reconstitution and Crystallization of Cas9 Endonuclease Bound to a Guide RNA and a DNA Target. Methods Enzymol 558, 515–537 (2015). 10.1016/bs.mie.2015.02.008

87 Hellman, L. M. & Fried, M. G. Electrophoretic mobility shift assay (EMSA) for detecting protein-nucleic acid interactions. Nat Protoc 2, 1849–1861 (2007). 10.1038/nprot.2007.249

88 Zhang, Y., Liu, T., Meyer, C. A., Eeckhoute, J., Johnson, D. S., Bernstein, B. E., Nusbaum, C., Myers, R. M., Brown, M., Li, W. & Liu, X. S. Model-based analysis of ChIP-Seq (MACS). Genome Biol 9, R137 (2008). 10.1186/gb-2008-9-9-r137

89 Liu, T. & Doherty, P. MACS: Model-based Analysis for ChIP-Seq, <https://macs3-project.github.io/MACS/index.html> (2025).

90 Amemiya, H. M., Kundaje, A. & Boyle, A. P. The ENCODE Blacklist: Identification of Problematic Regions of the Genome. Sci Rep 9, 9354 (2019). 10.1038/s41598-019-45839-z

91 Li, Q., Brown, J. B., Huang, H. & Bickel, P. J. Measuring reproducibility of high-throughput experiments. Annals of Applied Statistics 5, 1752–1779 (2011).

