## Supplemental_Figures for "Quantitative profiling of intrinsic dCas9-DNA recognition reveals key determinants of guide RNA performance"

[1]Department of Pediatrics, [2]Center for Advanced Genomic Technologies, Duke University, Durham, NC, USA, [3]Program in Cellular and Molecular Medicine, Boston Children's Hospital, Boston, MA, USA, [4]Department of Genomics and Computational Biology, University of Massachusetts Chan Medical School, Worcester, MA, USA, [5]Department of Cell Biology, [6]Department of Biomedical Engineering, [7]Program in Computational Biology and Bioinformatics, [8]Department of Biostatistics & Bioinformatics, Duke University, Durham, NC, USA, [9]Department of Systems Biology, University of Massachusetts Chan Medical School, Worcester, MA, USA

[†]These authors contributed equally for this work.


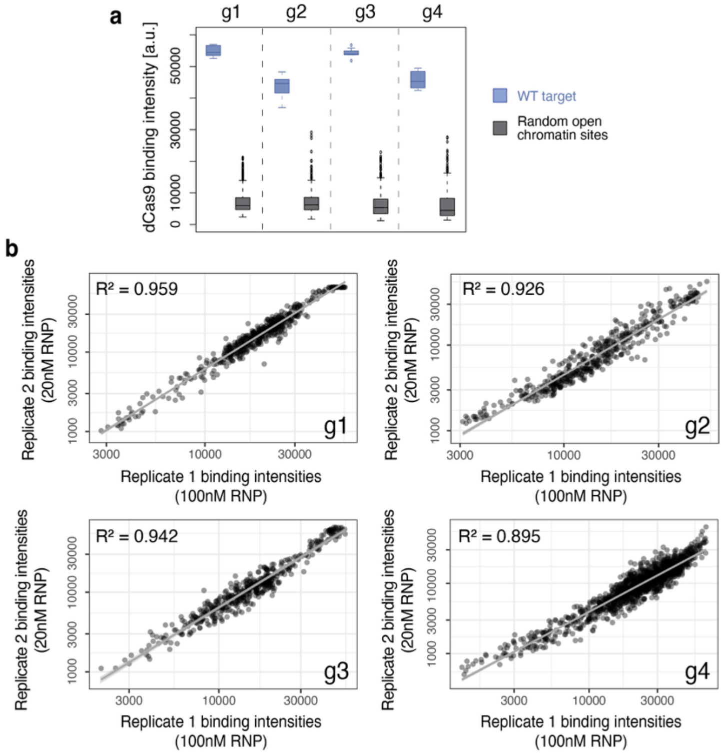


**Figure S1. TANGO measurements are gRNA-specific and reproducible across RNP concentrations and scan settings.** **a,** TANGO binding intensities for the four benchmark gRNAs (g1-g4), measured for WT target probes (blue) and random open-chromatin control probes (gray). **b,** Scatter plots comparing dCas9 binding intensities between two TANGO experiments performed at different RNP concentrations and scanner settings. Each panel shows data for a specific gRNA (g1, g2, g3, or g4) tested against its specific sub-library (**Methods**). X-axes show measurements collected at 100 nM RNP, using photomultiplier tube (PMT) voltage of 550 V and 15% laser power. Y-axes show measurements collected at 20 nM RNP, using PMT 600 V and 60% laser power. Each point represents a DNA probe from the corresponding guide sub-library, and lines indicate linear fits with $R^2$ shown on each panel, demonstrating high reproducibility across conditions. The guide-specific library for each guide includes the WT target probe, mutated target probes, and ChIP-identified off-targets from Kuscu et al.[1] (**Methods**).

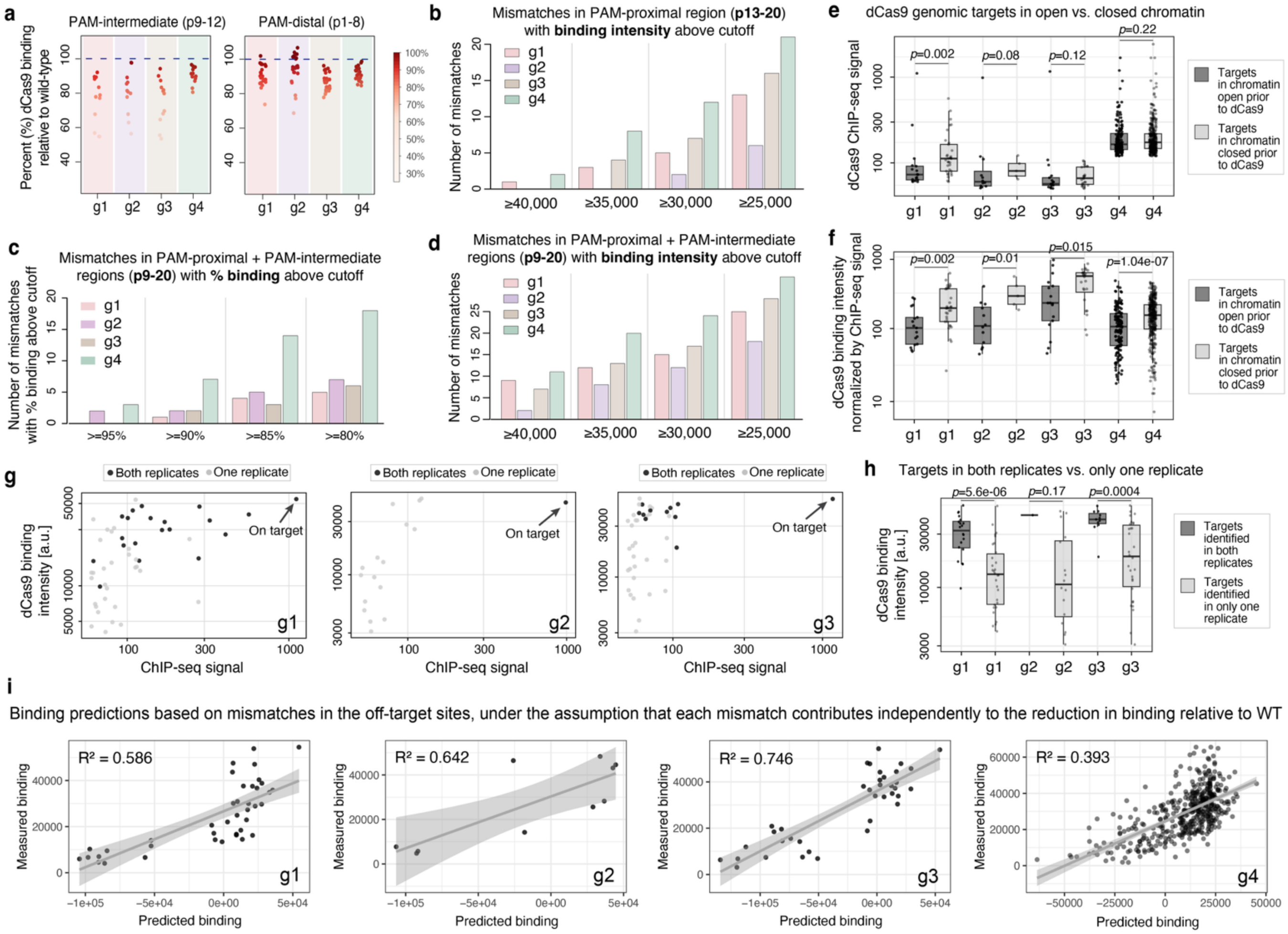


**Figure S2. TANGO-based mismatch analyses and ChIP-seq stratifications reinforce links between intrinsic binding and in-cell occupancy. a**, PAM-distal (positions 1-8) and PAM-intermediate (positions 9-12) mismatch tolerance summarized as percent binding for mismatched targets relative to the WT target (WT = 100%). Points are colored by %WT binding. Background colors are specific to each gRNA. **b**, For PAM-proximal mismatches (positions 13-20), plot shows the number of single-mismatch probes with TANGO binding intensity above the indicated cutoffs. **c**, For PAM-proximal + PAM-intermediate mismatches (positions 9-20), plot shows the number of single-mismatch probes retaining binding above the indicated %WT thresholds. **d**, For PAM-proximal + PAM-intermediate mismatches (positions 9-20), plot shows the number of single-mismatch probes with TANGO binding intensity above the indicated cutoffs. **e**, ChIP-seq signal for ChIP-identified genomic targets stratified by chromatin state (open vs. closed prior to dCas9) for each guide. P-values shown were computed using one-sided Mann-Whitney U tests. **f**, TANGO binding intensity normalized by ChIP-seq signal, for ChIP-identified genomic targets, computed as TANGO binding intensity divided by ChIP–seq signal, stratified by chromatin state (open vs. closed prior to dCas9). P-values shown were computed using one-sided Mann-Whitney U tests. **g**, TANGO binding intensity plotted against ChIP-seq signal for ChIP-identified genomic targets included on the TANGO array. Points are colored by whether the target was identified in both ChIP-seq replicates or in only one replicate. Arrows denote the on-target sites. Only g1-g3 are shown because g4 did not have two replicate ChIP-seq experiments in the study of Kuscu et al.[1]. **h**, TANGO binding intensities for targets identified in both ChIP-seq replicates vs. only one replicate, for each guide. P-values shown were computed using one-sided Mann-Whitney U tests. **i**, Scatter plots show a direct comparison between *in vitro* dCas9 binding at the ChIP-identified genomic off-targets (y-axis) vs. predicted *in vitro* binding based on the mismatch tolerance profiles of the four gRNAs. Predictions were made using a simple model where each mismatch in the off-target site contributes an independent penalty relative to the level of dCas9 binding to the perfectly matched on-target. Results show that this simple model is a good starting point for predicting dCas9 binding at putative off-targets sites, but it tends to over-penalize, likely because the penalties for consecutive mismatches are less than additive. Additional TANGO studies, using DNA libraries with a diversity of off-target sites, are needed in order to build more accurate predictive models.

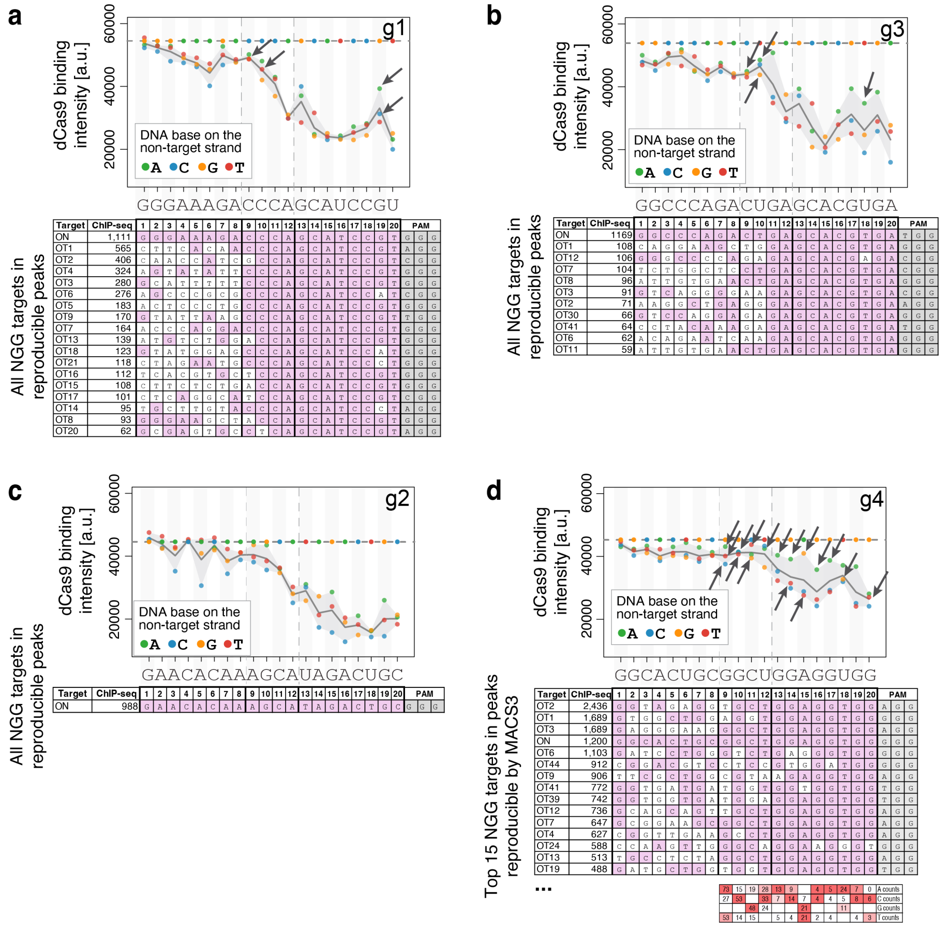


**Figure S3. Mismatches in the PAM-proximal and PAM-intermediate regions of ChIP-identified off-targets align with mismatches that retain strong binding in TANGO.** For each guide (g1-g4), the top panel shows the TANGO-derived mismatch tolerance profile across the target region (positions 1-20), with vertical dotted lines marking the PAM-distal, PAM-intermediate, and PAM-proximal regions. Points are individual single-mismatch probes (colored by the substituted base on the non-target strand), and the solid line shows the mean binding over the three mismatches at each position. Arrows mark the mismatches observed in ChIP-identified off-target sites (selected from reproducible dCas9 ChIP-seq peaks for g1-g3 and MACS3-reproducible peaks for g4; only off-targets with NGG PAMs are shown). The on-target and off-target sites are listed in the bottom panel for each guide, aligned to the gRNA spacer, with mismatched bases highlighted in white and the PAM sequence indicated on the right side. The ChIP-seq signals are shown on the left. For g4, which only has one replicate and thus could not be analyzed for reproducibility among replicates, we applied the ENCODE ChIP-seq pipeline[2], which uses the MACS3 algorithm[3] to call ChIP-seq peaks, and we focused this analysis on the peaks that were reproducible between the pipeline used by Kuscu et al.[1] and the ENCODE pipeline. For g4, the table shows the top 15 NGG target sites and a per-position nucleotide count summary across all off-target sites. Overall, these results show that the intrinsic mismatch tolerance of gRNAs, as captured by TANGO, contributes to their off-target binding in cells.

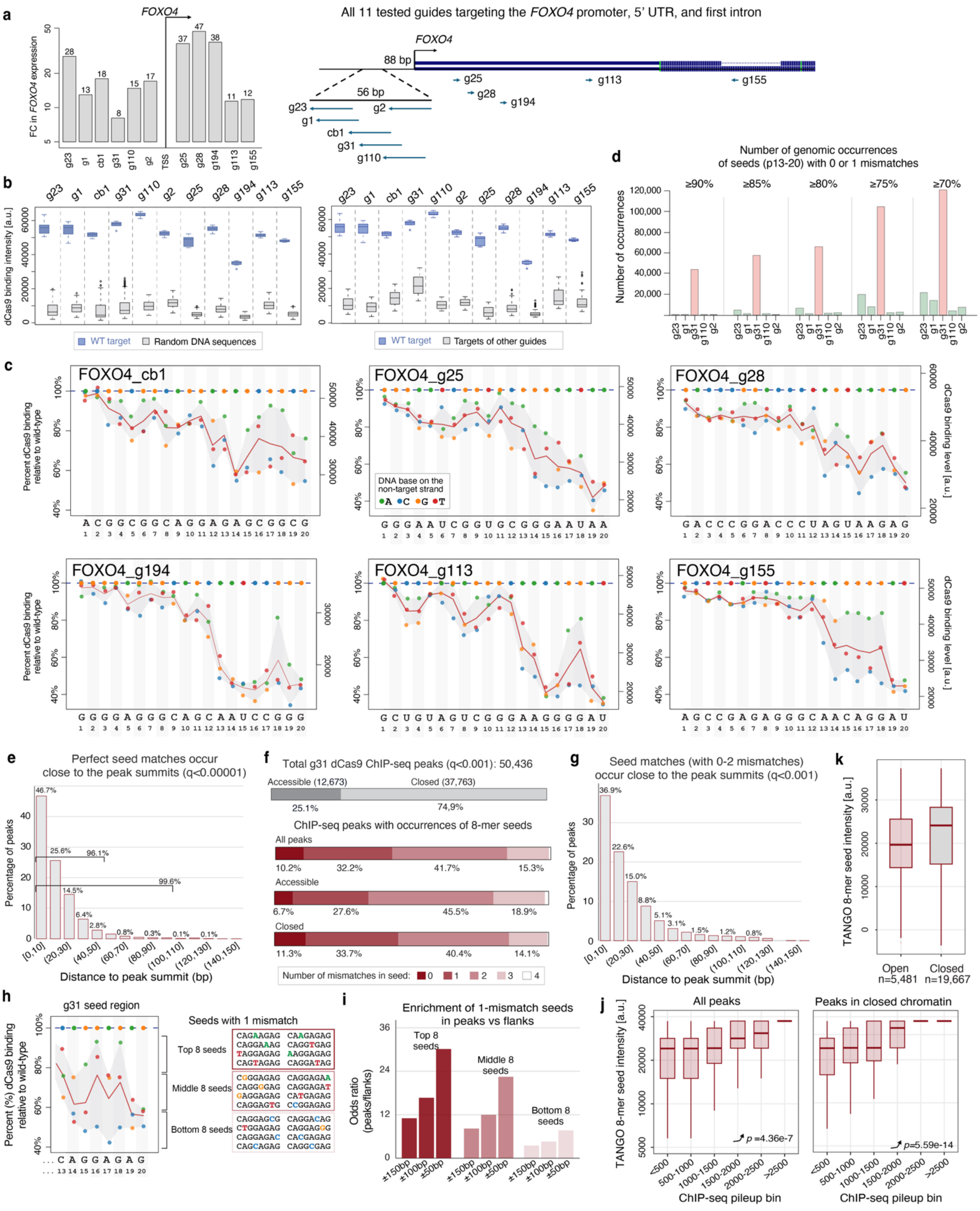


**Figure S4. TANGO binding and mismatch profiles for the *FOXO4* CRISPRa gRNA set. a**, CRISPRa activity in astrocyte cells, shown as fold change in *FOXO4* expression, for all tested 11 gRNAs targeting the *FOXO4* region[4]. **b**, TANGO data showing gRNA-specific binding to WT target probes compared to random DNA sequences (left) or targets of other gRNAs (right). **c**, Mismatch tolerance profiles for the six non-promoter *FOXO4* gRNAs, displayed as in Figure 3d-h. **d**, Number of occurrences, across the human genome (hg38), of 8-mer seeds (positions 13-20) with 0 or 1 mismatches that retained binding levels above the indicated thresholds relative to WT target site. Similar to Figure 3k, but including the perfect 8-mer seed. **e**, Occurrences of 8-mer seed sequences with 0 mismatches within ChIP-seq peaks, displayed as in Figure 4c. **f**, Number of ChIP-seq peaks identified for dCas9+g31 in astrocytes (**Methods**), at a less stringent q-value cutoff of 0.001, displayed as in Figure 4b. **g**, Occurrences of 8-mer seed sequences with 0-2 mismatches within ChIP-seq peaks, displayed as in Figure 4c. **h**, Single-mismatch seeds, i.e. 8-mers that are 1-nt different from the 8-bp PAM-proximal region of the perfect g31 target, grouped into 3 sets (high-affinity, medium-affinity, and low-affinity) based on their TANGO binding levels. **i**, Enrichment values, shown as odd-ratios of occurrences in the peaks vs. the flanks, for the three groups of seeds shown in panel h. **j**, Relationship between *in vitro* binding levels measured by TANGO (y-axis) vs. *in vivo* binding signals measured by ChIP-seq (x-axis), for the 25,148 g31 ChIP-seq peaks with a single occurrence of an 8-mer seed with 0-2 mismatches, displayed as in Figure 4g, but for either all peaks (left) or peaks in closed chromatin (right). P-values shown are according to Jonckheere-Terpstra tests for increasing trend. **k**, The same peaks as in panel j, but comparing the TANGO 8-mer seed intensities for peaks in open vs. closed chromatin prior to dCas9 recruitment (one-sided Mann-Whitney U test p=2.7e-67).

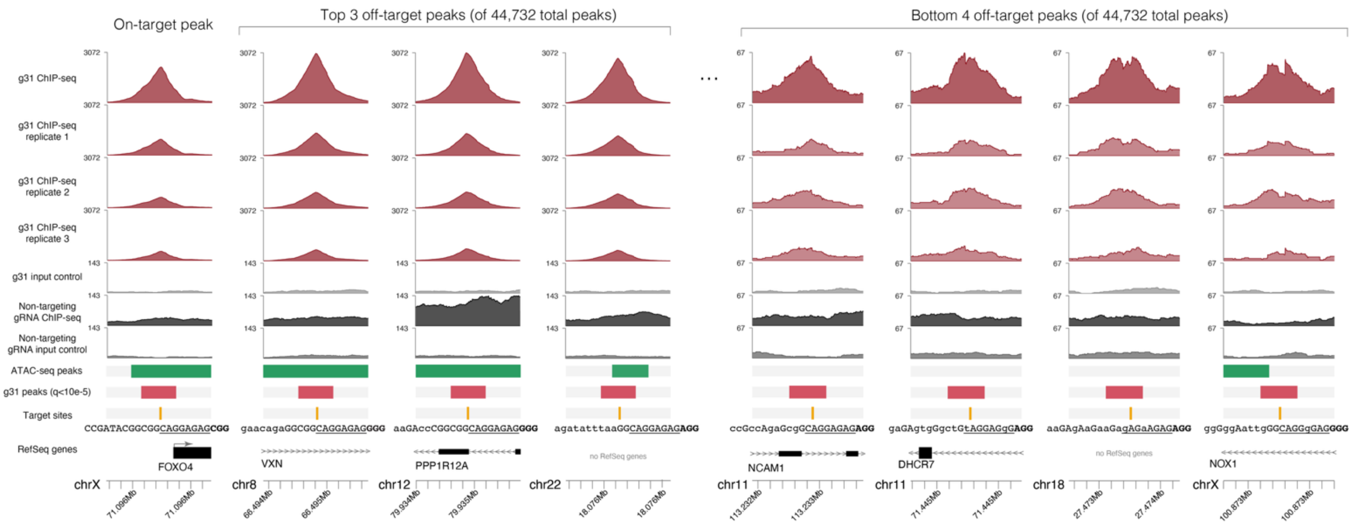


**Figure S5. Genome browser views of g31 RNP ChIP-seq peaks.** Left: the on-target peak in the *FOXO4* promoter shows high ChIP-seq signal, as expected. Middle: the top three g31 ChIP-seq peaks that have the highest read pileups. We note that the on-target peak is ranked 34th in the list of 44,732 peaks, so the peaks shown here have higher signal than the on-target peak. Right: the bottom four peaks that have the lowest read pileups among the 44,732 high-confidence, highly-reproducible g31 ChIP-seq peaks. Plots show that all peaks contain a match to the g31 target sequence followed by NGG, with at most 2 mismatches in the 8-mer seed region. The seed region is underlined, and lowercase letters are used to indicate mismatches compared to the perfect target.

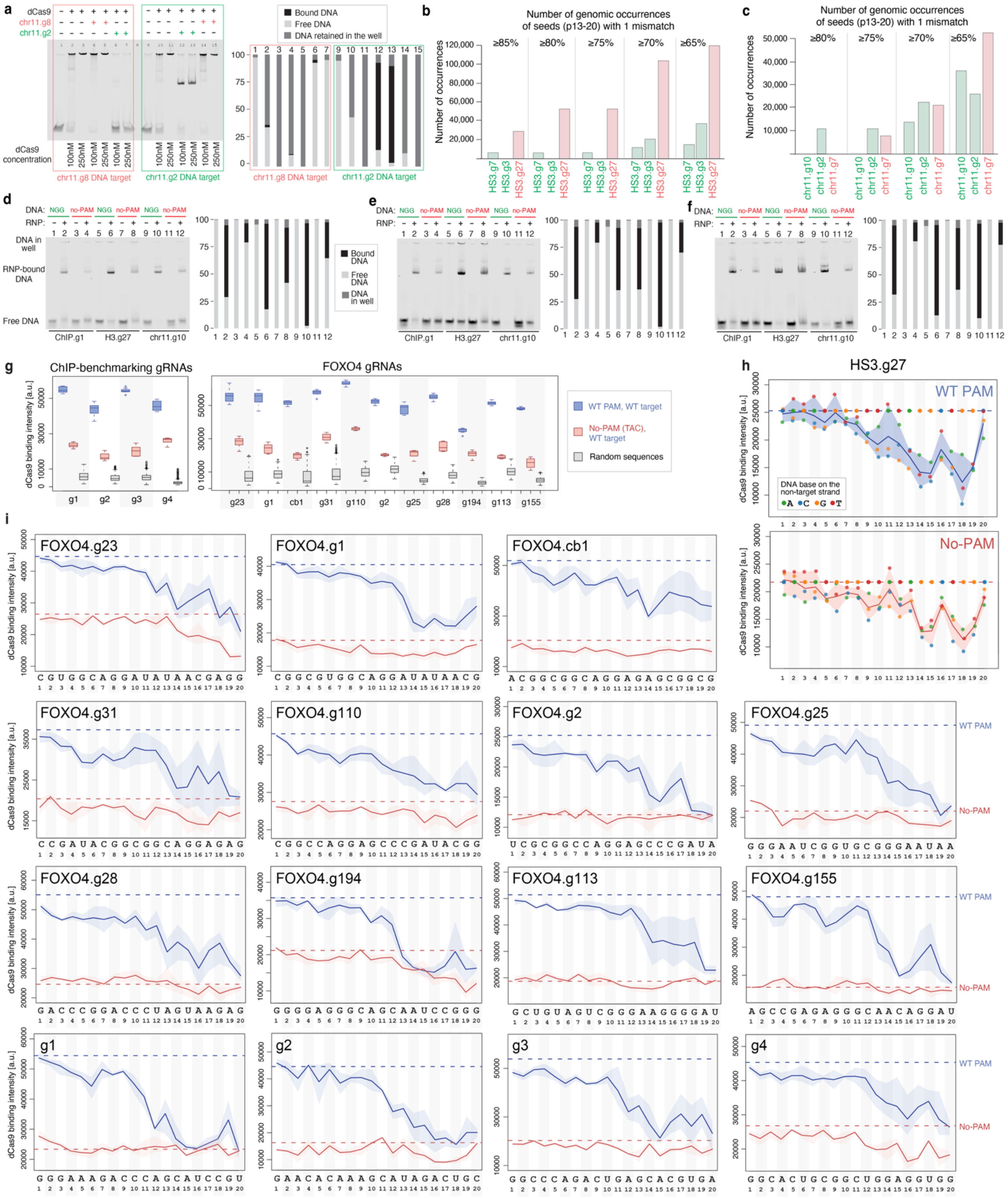


**Figure S6. TANGO reveals insights into the DNA-binding patterns of CRISPRi and CRISPRa gRNAs. a**, EMSA gels providing additional evidence that non-working gRNA chr11.g8 does not assemble efficiently with dCas9, as its gel pattern (with the DNA largely retained in the well) matches that of apo-dCas9. In contrast, working guide chr11.g2 shows the expected shift at its DNA target, and no shift at the DNA target of chr11.g8. **b**, Number of occurrences, across the human genome (hg38), of 8-mer seeds (positions 13-20) with a single mismatch and with binding levels above the indicated thresholds relative to WT, for gRNAs targeting the HS3 region of *HBE1*. **c,** Similar to a, but for gRNAs targeting the chr11.1735 region of *LMO2*.

Similarly to Figure 5, green indicates "working" gRNAs, while red indicates "non-working" gRNAs. **d-f**, EMSAs validating that dCas9 RNPs can bind well to perfectly complementary DNA targets even when they are adjacent to a non-PAM trinucleotide (here 5'-TAC-3'). EMSA gels were run with 5nM double-stranded DNA target and 250nM RNP, on 8% TBE gels, for 90min @100V, with 6ul per lane. EMSAs were run for HS3.g27 and chr11.10, the two outlier gRNAs that show the strongest NGG-independent binding, and with the g1 gRNA from the ChIP-benchmarking group, denoted here as ChIP.g1, which showed weak binding to no-PAM probes in TANGO. Panels e and f show EMSA where RNP:DNA binding was measured in the presence of crowders, 2mM spermidine and 4.5% Ficoll400, respectively. **g**, TANGO binding intensities for the four ChIP-benchmarking and the eleven *FOXO4* guides on WT PAM WT target probes (dark blue), alternative NGG PAM variants (light blue), no-PAM variants (red), and negative control targets (here, random DNA sequences; gray). **h,** Comparison between mismatch tolerance profiles for HS3.g27 at probes containing either the native NGG PAM (top) or the non-permissive no-PAM trinucleotide TAC (bottom). These profiles are simplified and shown as overlapping profiles in Figure 6c. **i**, Mismatch tolerance profiles four ChIP-benchmarking and the eleven *FOXO4* guides for probes containing either the native NGG PAM (blue) or the non-permissive no-PAM trinucleotide TAC (red). Colored lines show the mean binding intensity at each position, calculated over the three possible mismatches at that position. Horizontal dotted lines show the binding level at the WT target site with either the WT PAM (blue) or the no-PAM TAC (red). Blue and red shading shows the range of binding intensities across the mismatches at each position.

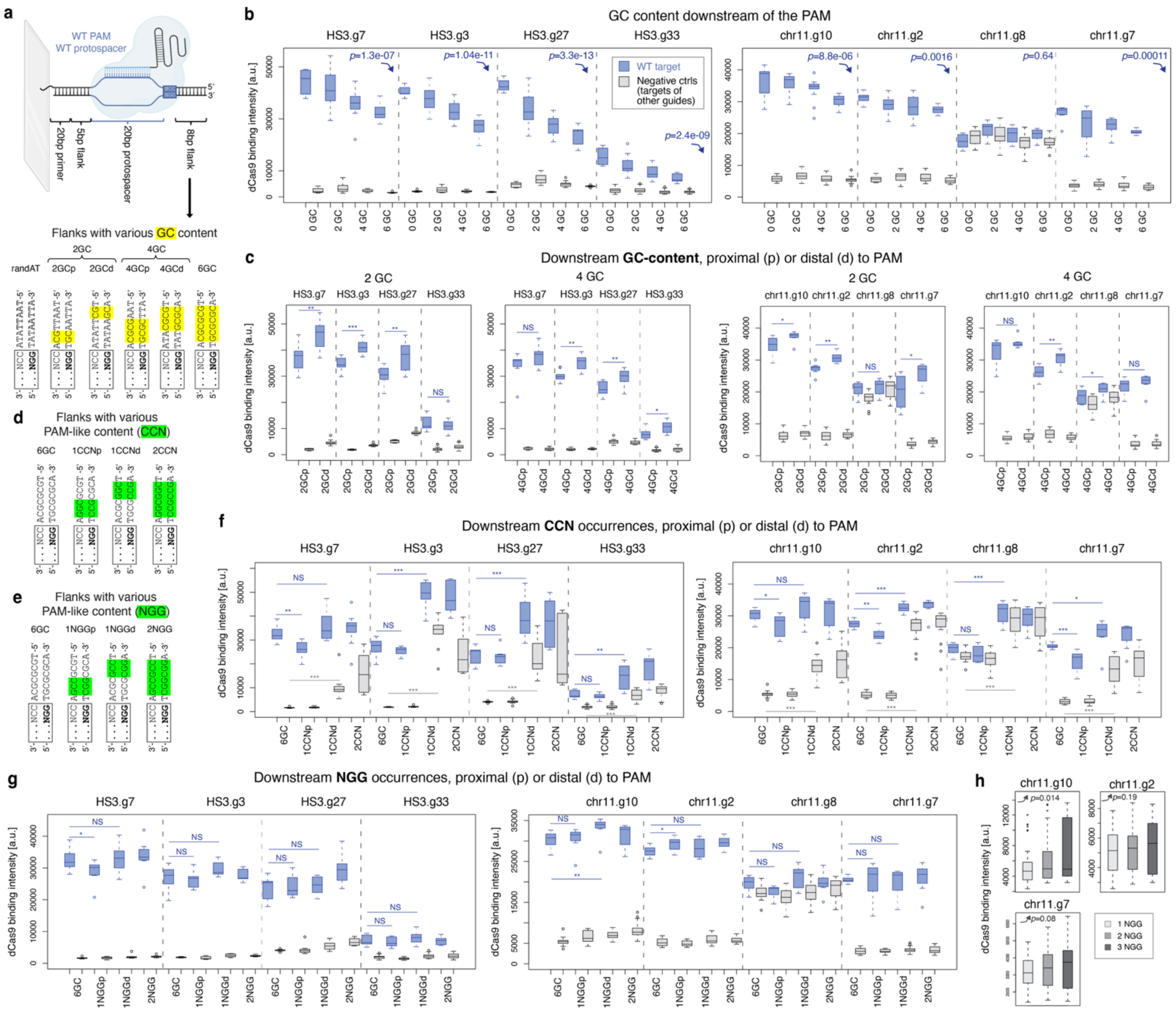

**Figure S7. The effects of downstream PAM-flanking sequence on dCas9 binding. a**, Top: probe design to test the effects of the 8-bp region immediately 3′ of the PAM and outside the protospacer, i.e. the downstream PAM-flanking sequence. The protospacer and WT NGG PAM regions were held constant, while the 8-bp flank immediately downstream the PAM (i.e. toward the free end of the 60-bp probe) was systematically varied. Bottom: downstream PAM-flanking sequence variants used in panels b and c. Starting from a 0GC backbone (8 A/T bases, 0 G/C bases), variants were generated with 2, 4, or 6 G/C bases (2GC, 4GC, 6GC) in the 8-bp flank. For the 2GC and 4GC variants, G/C bases were placed either PAM-proximal (2GCp, 4GCp) or PAM-distal (2GCd, 4GCd) within the flank. **b**, Effect of downstream PAM-flanking GC fraction on TANGO-measured dCas9 binding for HS3 (left) and chr11.1734 (right) gRNA sets using the designs in panel a. Box plots show binding to the WT targets (blue) and to negative controls (here, targets of other, non-overlapping guides printed on the same array; gray). P-values shown are according to Jonckheere-Terpstra tests for decreasing trend. **c**, Effect of G/C placement within the 8 bp downstream PAM flank, for fixed GC content (2GC and 4GC designs, as in a) on TANGO-measured binding for HS3 (left) and chr11.1734 (right) guide sets. Box plots are shown as in b. Statistical significance was assessed using one-sided Mann-Whitney U tests with the following p-value cutoffs: *** p<0.0005, ** p<0.005, * p<0.05, NS p≥0.05. **d,** Schematic of “extra CCN” designs constructed on the 6GC backbone: CCN-proximal (1CCNp), CCN-distal (1CCNd), and CCN×2 (proximal+distal, 2CCN). CCN corresponds to an NGG on the opposite strand relative to the WT PAM orientation. **e,** Schematic of “extra NGG” designs constructed on the 6GC backbone: NGG-proximal (1NGGp), NGG-distal (1NGGd), and NGG×2 (proximal+distal, 2NGG). **f,** TANGO binding intensities for the WT targets and negative controls for the CCN-series variants. Statistical significance was assessed using one-sided Mann-Whitney U tests with the following p-value cutoffs: *** p<0.0005, ** p<0.005, * p<0.05, NS p≥0.05. **g,** TANGO binding intensities for the WT targets and negative controls for the NGG-series. Statistical significance was assessed using one-sided Mann-Whitney U tests with the following p-value cutoffs: *** $p<0.0005$, ** $p<0.005$, * $p<0.05$, NS $p\geq0.05$. **h,** TANGO binding intensities for random DNA sequences with 1, 2, or 3 NGG motifs, which were included in the TANGO DNA library for the chr11.1734 guide set. Data is shown only for chr11.g10, chr11.g2, and chr11.g7, the three gRNAs in this set that have guide-specific DNA-binding activity. P-values shown are according to Jonckheere-Terpstra tests for increasing trend.

### REFERENCES


1 Kuscu, C., Arslan, S., Singh, R., Thorpe, J. & Adli, M. Genome-wide analysis reveals characteristics of off-target sites bound by the Cas9 endonuclease. *Nature biotechnology* **32**, 677-683 (2014). https://doi.org:10.1038/nbt.2916

2 Hitz, B. C., Jin-Wook, L., Jolanki, O., Kagda, M. S., Graham, K., Sud, P., Gabdank, I., Strattan, J. S., Sloan, C. A., Dreszer, T., Rowe, L. D., Podduturi, N. R., Malladi, V. S., Chan, E. T., Davidson, J. M., Ho, M., Miyasato, S., Simison, M., Tanaka, F., Luo, Y., Whaling, I., Hong, E. L., Lee, B. T., Sandstrom, R., Rynes, E., Nelson, J., Nishida, A., Ingersoll, A., Buckley, M., Frerker, M., Kim, D. S., Boley, N., Trout, D., Dobin, A., Rahmanian, S., Wyman, D., Balderrama-Gutierrez, G., Reese, F., Durand, N. C., Dudchenko, O., Weisz, D., Rao, S. S. P., Blackburn, A., Gkountaroulis, D., Sadr, M., Olshansky, M., Eliaz, Y., Nguyen, D., Bochkov, I., Shamim, M. S., Mahajan, R., Aiden, E., Gingeras, T., Heath, S., Hirst, M., Kent, W. J., Kundaje, A., Mortazavi, A., Wold, B. & Cherry, J. M. The ENCODE Uniform Analysis Pipelines. *bioRxiv* (2023). https://doi.org:10.1101/2023.04.04.535623

3 Liu, T. & Doherty, P. *MACS: Model-based Analysis for ChIP-Seq*, <https://macs3-project.github.io/MACS/index.html> (2025).

4 Reisman, S. J., Zhu, W., Miller, S. E., Halabi, D., Sangvai, N., Crawford, G. E., Gordan, R. & Gersbach, C. A. Mismatch tolerance of a gRNA for CRISPR-based gene activation confers broad activity critical for cell reprogramming. *bioRxiv* (2026). https://doi.org:10.64898/2026.02.01.703129
